# Lrp10 suppresses IL7R limiting CD8 T cell homeostatic expansion and anti-tumor immunity

**DOI:** 10.1101/2023.12.08.570738

**Authors:** Jamie Russell, Luming Chen, Aijie Liu, Jianhui Wang, Subarna Ghosh, Xue Zhong, Hexin Shi, Bruce Beutler, Evan Nair-Gill

## Abstract

Signals emanating from the T cell receptor (TCR), co-stimulatory receptors, and cytokine receptors each influence CD8 T cell fate. Understanding how these signals respond to homeostatic and microenvironmental cues can reveal new ways to therapeutically direct T cell function. Through forward genetic screening in mice, we discovered that loss-of-function mutations in *LDL receptor related protein 10* (*Lrp10*) caused naïve and central memory CD8 T cells to accumulate in peripheral lymphoid organs. *Lrp10* encodes a conserved cell surface protein of unknown immunological function. Lrp10 was induced with T cell activation and its expression post-translationally suppressed IL7 receptor (IL7R) levels. Accordingly, *Lrp10* deletion enhanced T cell homeostatic expansion through IL7R signaling. *Lrp10*-deficient mice were also intrinsically resistant to syngeneic tumors. This phenotype depended on dense tumor infiltration of CD8 T cells that displayed increased memory cell characteristics, reduced terminal exhaustion, and augmented responses to immune checkpoint inhibition. Here, we present Lrp10 as a new negative regulator of CD8 T cell homeostasis and a host factor that controls tumor resistance with implications for immunotherapy.

## INTRODUCTION

CD8 T cells circulate through peripheral lymphoid organs until encountering their cognate antigen. Antigen-induced activation leads to clonal expansion, expression of effector molecules, and migration to inflamed tissues. Clearance of the antigenic stimulus is followed by apoptosis of most antigen specific cells (D’Cruz *et al*, 2009). A small number of memory cells remain that are long-lived, variably capable of self-renewal, and can rapidly respond to subsequent challenges (Jameson & Masopust, 2018). When antigenic stimulation persists, for example within the tumor microenvironment (TME), or during a chronic viral infection, the transition from the effector phase to the memory phase is corrupted. In these instances, CD8 T cell function is impaired through a differentiation process known as “exhaustion” wherein subpopulations of tumor reactive clones with memory and stem-like features continuously propagate a terminally exhausted pool (Zehn *et al*, 2022). An array of transcription and epigenetic factors promote or oppose T cell exhaustion (Belk *et al*, 2022; Kaech & Cui, 2012). Additionally, signals transmitted through the T cell receptor (TCR), inhibitory and activating co-receptors, and cytokine receptors influence CD8 T cell fate according to extrinsic cues (Giles *et al*, 2023; Huang & August, 2015; Wei *et al*, 2019). How these externally derived signals integrate with cell-intrinsic gene regulatory programming remains uncertain. Therefore, identifying mechanisms that control the intensity and duration of externally derived signals could inform CD8 T cell-based immunotherapies.

To define new mechanisms that control T cell homeostasis and differentiation, we performed a forward genetic screen in randomly mutagenized mice that measured the proportions of T cells circulating in the peripheral blood. Here, we report that a gene called *LDL receptor Related Protein 10* (*Lrp10*) plays an important role in the homeostasis and differentiation of peripheral CD8 T cells through its effects on the interleukin 7 receptor (IL7R). *Lrp10* encodes a putative endocytic receptor that is a member of the LDL Receptor related Protein (LRP) superfamily (Sugiyama *et al*, 2000). Most LRPs bind and internalize diverse ligands through large extracellular domains that contain cysteine-rich LDL ligand-binding domains (LBD) and Epidermal Growth Factor (EGF) homology domains (Lane-Donovan *et al*, 2014). In contrast, Lrp10, along with Lrp3 and Lrp12, form a distinct subfamily of orphan LRPs with relatively small extracellular regions that contain LBDs and C1r/C1s, Uegf, Bmp1 (CUB) domains. Lrp10 was shown previously to facilitate intracellular vesicle trafficking in neuronal cells and astrocytes in cell culture (Brodeur *et al*, 2012). It also negatively regulated the growth of myeloid leukemia cells in mice (Ramakrishnan *et al*, 2020). Currently, there is little knowledge of its function *in vivo* and it has no previously described role in immune homeostasis. By deleting *Lrp10* in mice, we have discovered that Lrp10 prevents accumulation of naïve and memory CD8 T cells in secondary lymphoid organs, limits IL7R expression, suppresses T cell homeostatic expansion, and impairs anti-tumor immune responses.

## RESULTS

### Forward genetic screening reveals *Lrp10* is critical for normal CD8 T cell and NK cell homeostasis in mice

To identify new determinants of immune homeostasis, we screened the peripheral blood of mice mutagenized with N-ethyl-N-nitrosourea (ENU) with flow cytometry (Wang *et al*, 2015; Xu *et al*, 2021). Several mice from a single pedigree showed an increased proportion of CD8 T cells and a decreased proportion of natural killer (NK) cells, a phenotype that we named *chowmein*. Automated meiotic mapping linked the *chowmein* phenotype to a missense mutation in *LDL receptor related protein 10* (*Lrp10*) using a recessive model of inheritance (Fig. 1A and B).

**FIGURE 1:**
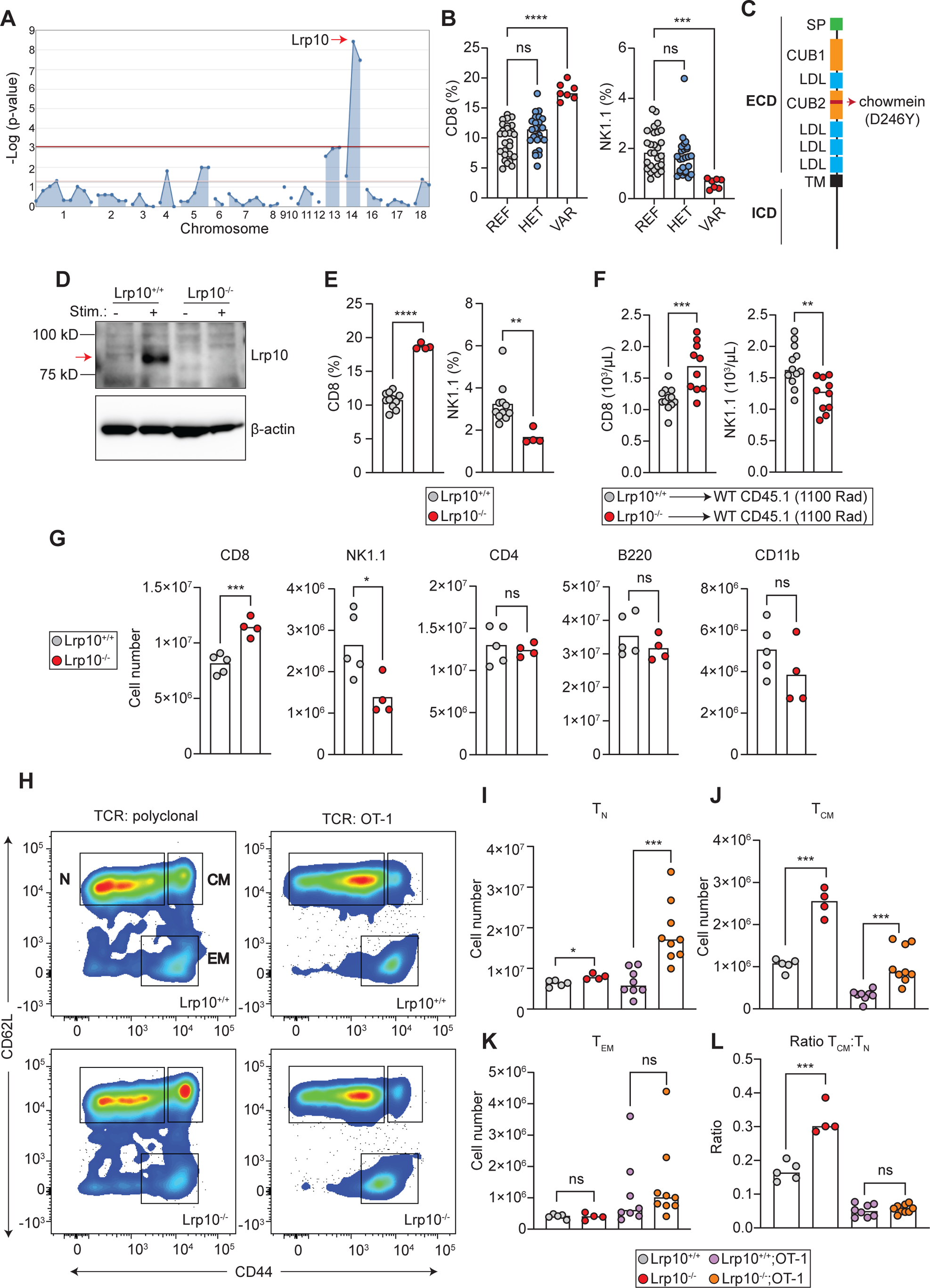
Lrp10 deletion increases CD8 T cells. **A:** Manhattan plot showing linkage between an ENU-induced point mutation in *Lrp10* and increased peripheral CD8 T cells. **B:** Frequency of peripheral CD8 T cells and NK cells in the *chowmein* pedigree. **C:** Domain structure of Lrp10 and location of the ENU-induced substitution. **D:** Expression of Lrp10 in resting and stimulated splenic CD8 T cells. **E:** Frequency of CD8 T and NK cells in the peripheral blood of CRISPR *Lrp10^−/−^* mice. **F:** Number of peripheral CD8 T cells and NK cells from lethally irradiated mice transplanted with *Lrp10^+/+^* or *Lrp10^−/−^* bone marrow. **G:** Numbers of the major splenic immune lineages. **H:** Representative FACS plot of CD8 T cell subpopulations from *Lrp10^+/+^* and *Lrp10^−/−^* mice harboring a polyclonal repertoire and a restricted repertoire (OT-1). **I - K:** Quantification of CD8 T cell subpopulations dependent on TCR repertoire. **L:** Ratio of T_CM_ : T_N_ cells. *P < 0.01, **P < 0.01, ***P < 0.001, ****P < 0.0001 by one-way ANOVA with Dunnett’s multiple comparisons test (A), Mann-Whitney test (B, J, K, L), or two-tailed unpaired *t* test (E, F, G, I, J). Results are representative of three (D) and two (F) independent experiments.

Lrp10 possesses two CUB ligand binding domains interspersed with LBDs, a single-pass transmembrane region, and a proline-rich intracellular region (Fig. 1C). The *chowmein* allele encoded an aspartate to tyrosine substitution at position 246 (D246Y) in the second CUB domain of the extracellular region. Lrp10 was expressed at low levels in unstimulated CD8 T cells and its expression increased with T cell activation (Fig. 1D).

To verify that the increase in CD8 T cells observed was caused by the loss of Lrp10, we used CRISPR-Cas9 to create a constitutive knock-out allele of *Lrp10*. *Lrp10* knock-out mice (*Lrp10^−/−^*) were born at the expected Mendelian ratio, appeared outwardly normal, and were viable and fertile. *Lrp10^−/−^* mice showed an increased proportion of CD8 T cells and a reduction in NK cells in the peripheral blood, confirming that loss of Lrp10 function was responsible for the *chowmein* phenotype (Fig. 1E). Lethally irradiated *Rag2^−/−^* mice reconstituted with *Lrp10^−/−^*bone marrow had increased numbers of peripheral CD8 T cells and reduced numbers of NK cells compared to those receiving *Lrp10^+/+^* bone marrow, indicating that these phenotypes were hematopoietic-intrinsic (Fig. 1F).

### *Lrp10^−/−^* mice accumulate naïve and central memory CD8 T cells in a TCR repertoire-dependent manner

Spleens from *Lrp10^−/−^* mice harbored increased absolute numbers of CD8 T cells and reductions in NK cell numbers reflecting what we observed in the peripheral blood (Fig. 1G). There was no difference in the numbers of CD4+ T cells, B220+ B cells, or CD11b+ myeloid cells between *Lrp10^+/+^*and *Lrp10^−/−^* mice.

We first considered that the increase in peripheral CD8 T cells in *Lrp10^−/−^* mice might be due to preferential skewing toward the CD8+ lineage during thymic selection. However, thymic cellularity was the same between *Lrp10^+/+^* and *Lrp10^−/−^* mice with respect to double negative, double positive, and CD4 and CD8 single positive thymocytes (Fig. S1A).

The peripheral CD8 T cell population in mice contains three broad subpopulations defined by expression of the lymph node homing receptor CD62L and the tissue homing receptor CD44: naïve cells that have not been exposed to antigen (T_N_, CD62L+CD44-), central memory cells (T_CM_, CD62L+CD44+) that circulate through secondary lymphoid organs, and effector memory cells that patrol peripheral tissues and exhibit lower levels of lymphoid recirculation (T_EM_, CD62L-CD44+) (Nolz *et al*, 2011). *Lrp10^−/−^* mice showed a slight increase in the number of CD8 T_N_ cells compared to *Lrp10^+/+^* mice (Fig. 1H-L). Interestingly, *Lrp10^−/−^* mice exhibited a 2-3-fold expansion in the CD8 T_CM_ population. In contrast, the number of CD8 T_EM_ cells was the same between *Lrp10^+/+^* and *Lrp10^−/−^*spleens. We did not observe any differences in the distribution of the memory subpopulations in *Lrp10^−/−^* CD4 T cells (Fig. S1B).

CD8 T_CM_ cells appearing in unimmunized mice can arise from endogenous/self-antigen exposure and homeostatic expansion driven by cytokines (Fry & Mackall, 2005; Jameson & Masopust, 2018; White *et al*, 2017). To define the role of TCR specificity in the accumulation of *Lrp10^−/−^*CD8 T_CM_ cells, we crossed *Lrp10^−/−^* mice to the OT-1 TCR transgenic strain in which the majority of CD8 T cells express a TCR specific for ovalbumin (ova). Compared to *Lrp10^−/−^* mice with a diverse TCR repertoire, *Lrp10^−/−^;OT-1* mice had a higher number of T_N_ and a lower number of T_CM_ cells (Fig. 1H-L). The ratio of T_CM_ to T_N_ cells was normalized in *Lrp10^−/−^;OT-1* compared to *Lrp10^−/−^* mice. Restricting the TCR repertoire did not change the number of CD8 T_EM_ cells in the spleen. These results show that restricting the CD8 TCR repertoire imparts a block in the conversion of *Lrp10^−/−^* T_N_ cells to T_CM_ cells and suggests that TCR responsiveness to endogenous/self-antigens is important for the accumulation of T_CM_ cells in *Lrp10^−/−^* mice.

### Normal antigen-specific proliferation and CD8 cytotoxic responses in *Lrp10^−/−^* CD8 T cells

We first hypothesized that *Lrp10* deletion sensitized TCR signaling to antigens that were present in low amounts or that had low TCR affinities. However, *Lrp10^+/+^;OT-1* and *Lrp10^−/−^;OT-1* cells showed similar proliferative responses to high and low doses of ova *in vitro* (Fig. S2A). Additionally, adoptively transferred *Lrp10^+/+^;OT-1* and *Lrp10^−/−^;OT-1* cells showed similar proliferative responses *in vivo* after immunization with SIINFEKL (a high affinity peptide antigen for the OT-1 TCR) or SIITFEKL (a peptide antigen with ∼100-fold lower TCR affinity).

We next assessed how loss of Lrp10 affected cytotoxic responses. *Lrp10^+/+^*and *Lrp10^−/−^* mice showed similar levels of killing SIINFEKL-pulsed target cells after immunization with ova and alum adjuvant (Fig. S2C). Consistent with lower circulating NK cell numbers, *Lrp10^−/−^*mice had a mild defect in killing MHC-Class I-deficient target cells (Fig. S2D).

Surprisingly, despite harboring increased numbers of T_CM_ CD8 cells apparently derived from self/endogenous antigen exposures, *Lrp10^−/−^*mice did not show spontaneous autoimmunity. We performed CD8 cytotoxicity assays at timepoints after ova-alum immunization and found that *Lrp10* deletion did not impart higher levels of target cell killing over time (Fig. S2E). Moreover, both *Lrp10^+/+^* and *Lrp10^−/−^*mice responded similarly to a boost with ova performed 90 days after immunization. Overall, these data show that although *Lrp10* deletion promotes the differentiation and accumulation of CD8 T_CM_ cells, it does not enhance TCR sensitivity or promote unrestrained CD8 T cell cytotoxicity or cytotoxic memory.

### *Lrp10* limits IL7 expression and T cell homeostatic proliferation

CD8 T_N_ and T_CM_ require IL7R signaling for differentiation and survival (Carrette & Surh, 2012; Schluns *et al*, 2000). Based on the accumulation of T_N_ and T_CM_ in *Lrp10^−/−^*mice, we next analyzed cell surface expression of IL7R (Fig. 2A). CD8 T_N_ and T_CM_ from *Lrp10^−/−^* mice showed ∼30% and ∼50% increased cell surface IL7R expression, respectively. CD8 T_EM_ cells displayed a bimodal expression pattern for IL7R. *Lrp10^−/−^* CD8 T_EM_ cells showed ∼35% increased cell surface IL7R within the IL7R+ subpopulation. More IL7R was also present on the surface *Lrp10^−/−^*CD4 T cell subsets, although not to the levels seen on *Lrp10^−/−^*CD8 T cells (Fig. S3A).

**FIGURE 2:**
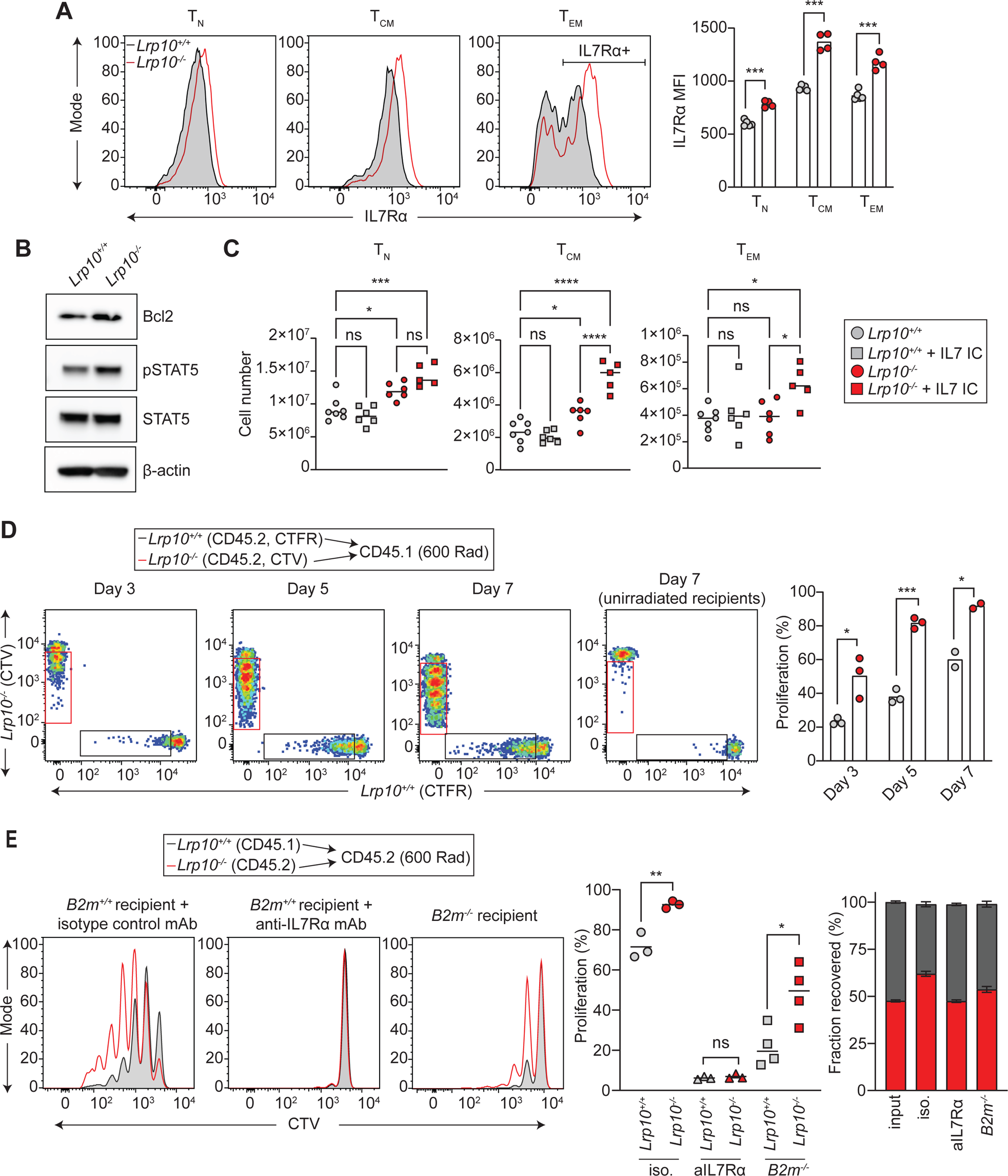
*Lrp10* deletion increases IL7R expression and function. **A:** IL7R expression on splenic CD8 T cell subpopulations. **B:** Expression of markers for IL7R signaling in resting splenic CD8 T cells. Results are representative of two independent experiments. **C:** Effect of IL7/anti-IL7 immune complex (IC) administration on splenic CD8 T cell populations. Data is combined from two-independent experiments with at least two mice per treatment group. **D:** Homeostatic expansion of differentially labeled naïve *Lrp10^+/+^*(CTFR) and *Lrp10^−/−^* (CTV) CD8 T cells in lymphopenic hosts. Proliferation was determined from the percentage of cells that underwent at least one cell division. Data is representative of three independent experiments. **E:** Homeostatic expansion of naïve *Lrp10^+/+^* (CD45.1) and *Lrp10^−/−^* (CD45.2) CD8 T cells injected into sub-lethally irradiated *B2m^+/+^* or *B2m^−/−^* (CD45.2) recipients. Mice were treated with anti-IL7R or isotype control antibodies as indicated. Homeostatic competitiveness was assessed based on the fraction of *Lrp10^+/+^*(gray bar) or *Lrp10^−/−^* (red bar) cells recovered on D7. Data is combined from one IL7R blockade experiment and one *B2m^−/−^*transplantation experiment. *P < 0.01, **P < 0.01, ***P < 0.001, ****P < 0.0001 by two-tailed unpaired *t* test (A, D, E) or one-way ANOVA with Tukey’s multiple comparisons test (C).

IL7R signaling is critical for thymic T cell development where it is expressed initially at the DN stage, turned off at the DP stage, and again expressed during selection of single-positive CD4 and CD8 T cells (Singer *et al*, 2008). There was no difference in IL7R expression in *Lrp10^−/−^*DN, DP, or CD4 single positive T cells (Fig. S3B). *Lrp10^−/−^* single positive CD8 T cells showed a small increase in IL7R expression compared to *Lrp10^+/+^*cells, although this difference was much less compared to peripheral CD8 T cells.

IL7 binds IL7R to activate STAT5, upregulate Bcl2, and promote T cell survival (Mazzucchelli & Durum, 2007; Rochman *et al*, 2009). Splenic *Lrp10^−/−^* CD8 T cells showed increased phosphorylated STAT5 and Bcl2 expression, consistent with enhanced basal IL7R signaling (Fig. 2B). Repeated injections of IL7/anti-IL7 immune complexes (IL7 IC) into mice has been shown to stimulate IL7R to increase T cell numbers (Boyman *et al*, 2008). We found that a single injection of IL7 IC caused preferential expansion of T_CM_ and T_EM_ in *Lrp10^−/−^*spleens (Fig. 2C), indicating that *Lrp10* deletion sensitized these populations to exogenous IL7.

IL7R signaling promotes T cell homeostatic proliferation during lymphopenia (Kimura *et al*, 2013; Tan *et al*, 2001). To further examine IL7 responsiveness in *Lrp10^−/−^* CD8 T cells, we tested their ability to proliferate under lymphopenic conditions. Naïve splenic CD8 T cells were harvested from *Lrp10^+/+^* and *Lrp10^−/−^* mice, differentially labeled with cell proliferation dye, and transplanted in equal numbers into syngeneic sub-lethally irradiated recipients (Fig. 2D). *Lrp10^−/−^*CD8 T cells showed a rapid increase in cell proliferation starting three days post-transfer. By 7 days, ∼90% of cells had undergone at least one round of cell division. In contrast, *Lrp10^+/+^* CD8 T cells showed lower rates of homeostatic expansion at early timepoints and by day 7 ∼60% of cells had undergone at least one round of division. Cells transferred into lymphocyte replete recipients showed no dye dilution, indicating that *Lrp10^−/−^*CD8 T cells did not spontaneously proliferate.

CD4 T cells are present in normal numbers in *Lrp10^−/−^*mice and display slightly increased levels of IL7R. Consistent with these findings, *Lrp10^−/−^* CD4 T cells showed mildly enhanced homeostatic proliferation upon transfer into sub-lethally irradiated recipients (Fig. S3C).

IL7R signaling and TCR signaling arising from self-peptide/MHC interactions each control T cell homeostatic proliferation (Kawabe *et al*, 2021). We next dissected how these distinct signals contributed to *Lrp10^−/−^*CD8 T cell homeostatic expansion. We labeled *Lrp10^+/+^* (CD45.1) and *Lrp10^−/−^* (CD45.2) naïve CD8 T cells with cell proliferation dye and adoptively transferred them into sub-lethally irradiated syngeneic mice. Recipient mice were then injected with an IL7R blocking antibody or an isotype control antibody (Fig. 2E). *Lrp10^−/−^* CD8 T cells from mice injected with the isotype control antibody showed the typical increased expansion phenotype. Conversely, anti-IL7R completely blocked proliferation of *Lrp10^+/+^* and *Lrp10^−/−^* CD8 T cells. *Lrp10^−/−^* CD8 T cells were recovered at a higher frequency from isotype control-injected mice and this competitive advantage was neutralized through IL7R blockade.

To test the effect of TCR signaling on CD8 T cell homeostatic expansion, we adoptively transferred *Lrp10^+/+^* (CD45.1) and *Lrp10^−/−^*(CD45.2) naïve CD8 T cells into sub-lethally irradiated syngeneic mice that lacked MHC-I expression (*Beta-2 microglobulin* knockout, *B2m^−/−^*). Transfer of CD8 T cells into *B2m^−/−^* mice substantially limited the homeostatic expansion of both *Lrp10^+/+^* and *Lrp10^−/−^*CD8 T cells (Fig. 2E). However, *Lrp10^−/−^* CD8 T cells continued to display higher levels of dye dilution and were recovered at increased frequencies. These findings show that increased homeostatic expansion of *Lrp10^−/−^*CD8 T cells depends entirely on IL7R signaling. While TCR and IL7R signaling combine to augment homeostatic expansion of *Lrp10^−/−^* CD8 T cells, they can proliferate at a reduced capacity in the absence of TCR-MHC-I interactions.

### Lrp10 post-translationally suppresses IL7R maturation

We next investigated how Lrp10 modulated IL7R expression. *IL7R* mRNA levels were the same in *Lrp10^+/+^* and *Lrp10^−/−^*CD8 T cells, indicating that Lrp10 reduced IL7R cell surface expression through a post-transcriptional mechanism (Fig. 3A). We used retroviruses encoding Lrp10, or an empty vector control, to complement *Lrp10^−/−^* CD8 T cells (Fig. 3B). Re-introducing Lrp10 reduced cell-surface IL7R on *Lrp10^−/−^* CD8 T cells, suggesting a direct link between Lrp10 and IL7R protein expression.

**FIGURE 3:**
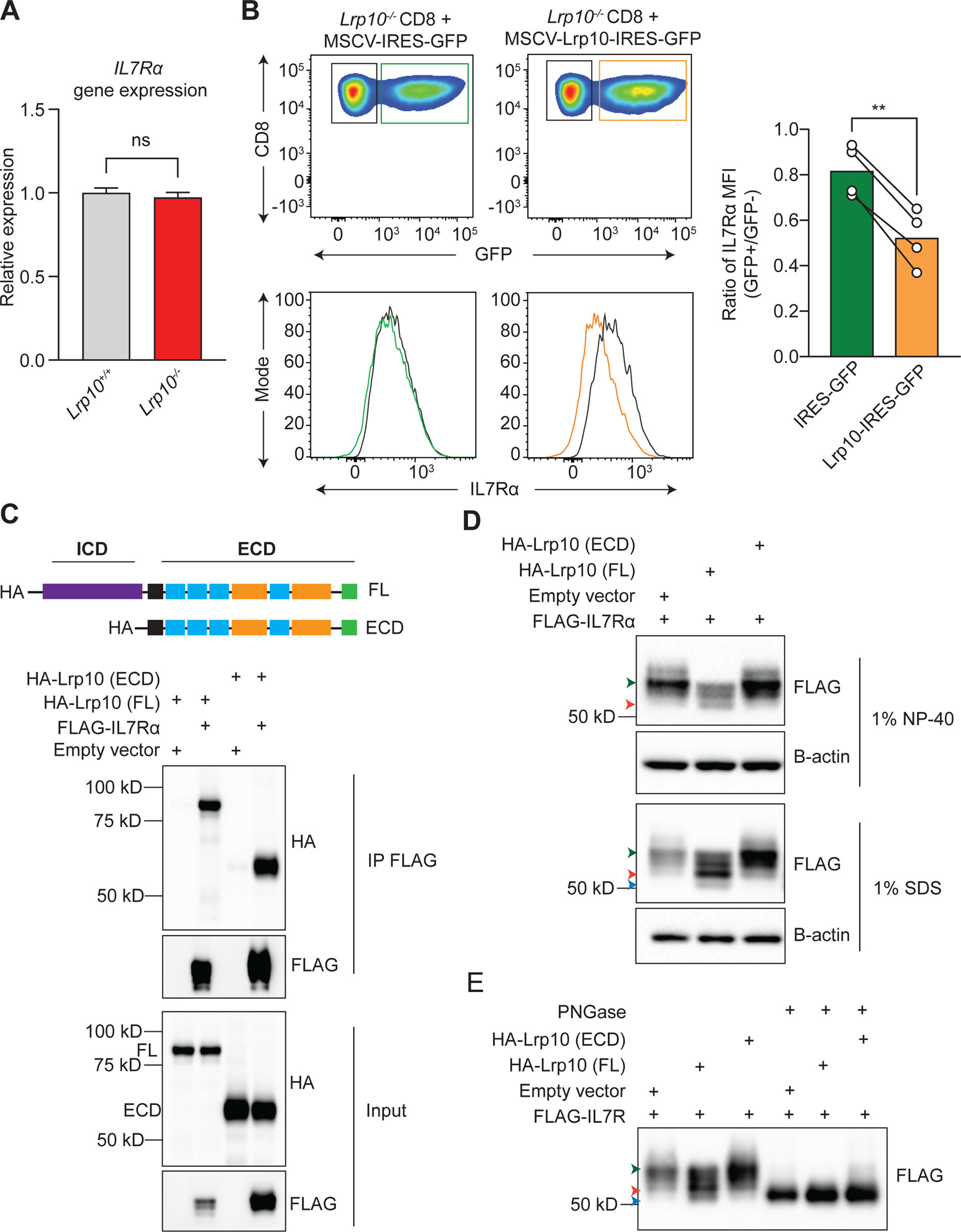
Lrp10 binds to IL7Rα and impairs its glycosylation. **A:** RT-qPCR measurement of *IL7Rα* gene expression in splenic CD8 T cells. Data presented as the mean and SD of three biological replicates. **B:** Representative FACS plots of cell surface IL7R expression on activated *Lrp10^−/−^* CD8 T cells infected with retroviruses encoding GFP-only (green trace) or Lrp10^WT^-IRES-GFP (yellow trace). IL7R levels on GFP+ cells were normalized to levels on the GFP-population (black trace). Bar graphs show results from four independent transduction experiments. **P < 0.01 by a two-tailed paired *t* test. **C:** Anti-FLAG co-IP of FLAG-IL7Rα with full length HA-Lrp10 (FL) or HA-Lrp10 extracellular domain (ECD) from transfected HEK 293T cells. **D:** FLAG-IL7Rα expression in transfected HEK 293T cells in the context of empty vector, HA-Lrp10 (FL), or HA-Lrp10 (ECD) under different lysis conditions. Colored arrows highlight the lower molecular weight IL7Rα isoforms. **E:** IP of FLAG-IL7Rα from transfected HEK 293T cells under denaturing conditions in the context of empty vector, HA-Lrp10 (FL), or HA-Lrp10 (ECD) followed by de-glycosylation with PNGase-F.

We hypothesized that Lrp10 may bind to IL7R to limit its expression. Heterologous expression of epitope tagged Lrp10 and IL7R in HEK 293T cells followed by co-immunoprecipitation (co-IP) of IL7R showed an interaction between these two proteins (Fig. 3C). This interaction was maintained after deletion of the Lrp10 ICD, indicating that IL7R bound to Lrp10 through the Lrp10 ECD.

In these co-IP experiments, we noted reduced levels of IL7R when it was co-expressed with full-length Lrp10, but not Lrp10-ECD (Fig. 3C). This was accompanied by IL7R isoforms that migrated at a lower molecular weight. To examine this phenomenon further, we expressed IL7R alone, with full-length Lrp10, or with Lrp10-ECD in HEK 293T cells. Upon extraction of cells with non-ionic detergent (1% NP-40) followed by SDS-PAGE analysis, we noted that full-length Lrp10 selectively decreased IL7R expression and induced lower molecular weight IL7R isoforms (Fig. 3D). Upon extraction of cells in a strong ionic detergent (1% SDS), we observed higher protein levels of IL7R in the context of full-length Lrp10 and more clearly resolved lower molecular weight IL7R isoforms that were not detected when IL7R was expressed alone or with Lrp10-ECD.

IL7R expressed alone, or with Lrp10-ECD, migrated at ∼60 kilodaltons (kD). The expected molecular weight of IL7R was ∼52 kD, which corresponded to the lowest molecular weight observed during co-expression with full-length Lrp10. It is know that IL7R is highly glycosylated, which positively affects its ability to bind IL7 (McElroy *et al*, 2009). We speculated that the lower molecular weight isoforms represented differentially glycosylated IL7R. Indeed, after treatment with a general deglycosylase (PNGase), all the observed isoforms of IL7R migrated at the predicted 52 kD molecular weight (Fig. 3E). Together, our data suggest a model where Lrp10 binds to IL7R through its ECD and uses its ICD to impair IL7R glycosylation and maturation to a fully functional cytokine receptor.

### Lrp10 limits CD8 T cell tumor infiltration and anti-tumor immunity

Tumor infiltration by cytotoxic CD8 T cells correlates with improved survival and positive responses to immune checkpoint inhibition (Galon & Bruni, 2019; Lee & Ruppin, 2019; Li *et al*, 2021). Our data thus far showed that *Lrp10^−/−^* mice harbored increased numbers of CD8 T cells that unexpectedly did not exhibit superior cytotoxic activity or recall responses after immunization with a model antigen. Therefore, we asked how *Lrp10* deletion might affect anti-tumor immune responses. We subcutaneously inoculated *Lrp10^+/+^*and *Lrp10^−/−^* mice with MC38 cells which give rise to syngeneic, highly immunogenic tumors derived from a chemically induced murine colon cancer. *Lrp10^−/−^* mice showed enhanced resistance to MC38 tumor growth (Fig. 4A and B). Immune phenotyping of infiltrating cell populations revealed that tumors from *Lrp10^−/−^*mice harbored increased numbers of CD8 T cells, but similar numbers of CD4 T cells and macrophages (Fig. 4C). Within the CD4 population, the frequency of regulatory T cells (T_reg_) was the same between each strain (Fig. S4A and B). Although *Lrp10^−/−^* mice have fewer circulating NK cells, the numbers of NK1.1+ cells were similar in tumors from *Lrp10^+/+^* and *Lrp10^−/−^* mice (Fig. 4C). Notably, the tumor resistance phenotype in *Lrp10^−/−^* mice was limited to the highly immunogenic MC38 tumor. We did not see a difference in tumor growth rate, or CD8 T cell infiltration, between *Lrp10^+/+^* and *Lrp10^−/−^* mice inoculated with the “immunologically cold” B16-F10 melanoma cell line (Fig. S4C and D).

**FIGURE 4:**
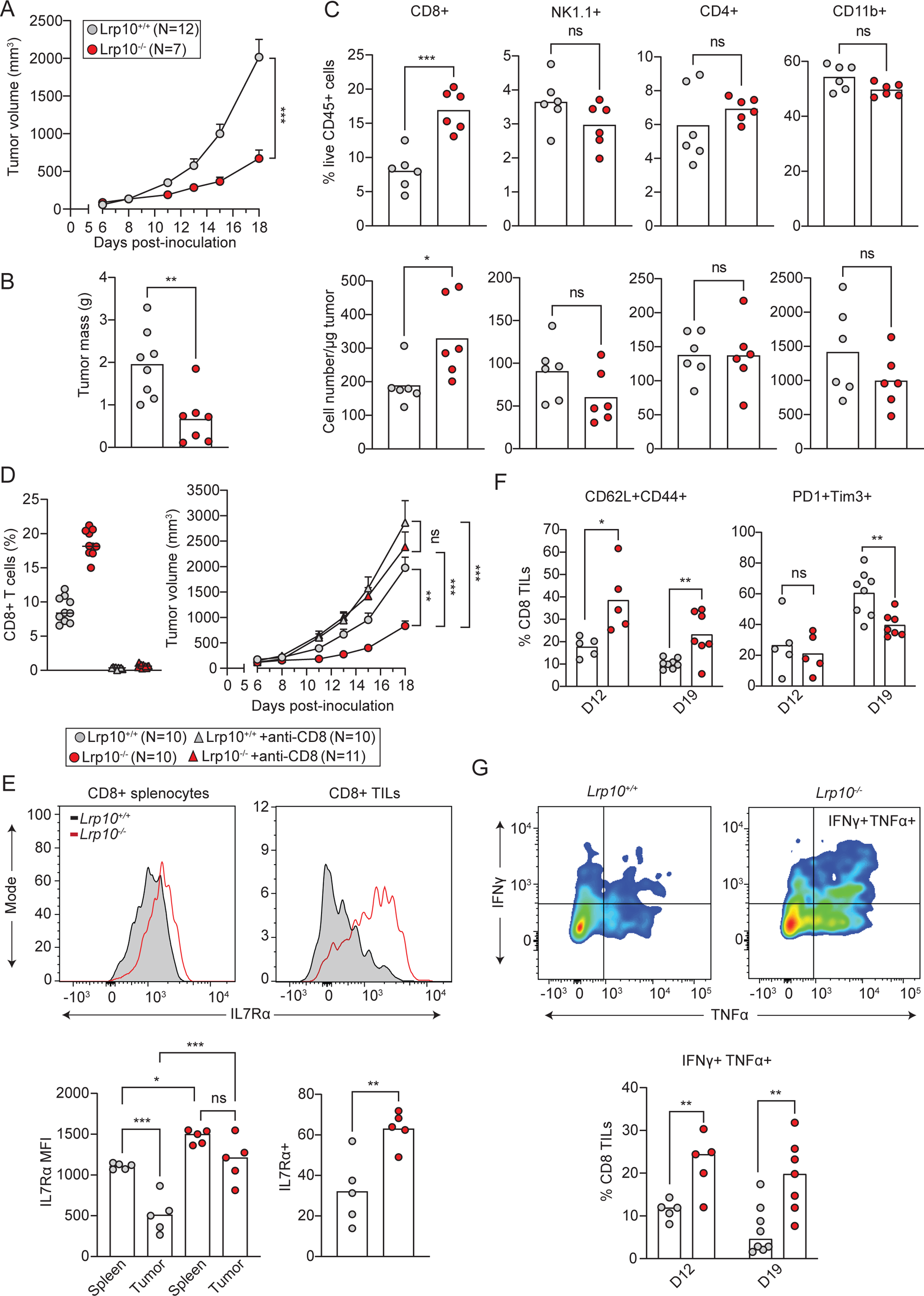
*Lrp10* deletion imparts tumor resistance. **A:** MC38 tumor volumes. Data is presented as mean and SEM. Differences in tumor volume were compared on D18. **B:** D18 MC38 tumor mass. **C:** Quantification of CD45+ immune cells in D18 MC38 tumors. **D:** Effect of CD8 T cell depletion on MC38 tumor growth. Data presented as mean and SEM. Differences in tumor volume were compared on D18. **E:** FACS plot of IL7R expression on spleen and tumor CD8 T cells from D12 MC38 tumors, quantification of IL7Rα MFI, and frequency of IL7Rα+ CD8 TILs. **F:** Frequency of memory phenotype (CD62L+CD44+) and terminally exhausted (PD1+Tim3+) CD8 TILs in MC38 tumors at the indicated time points. **G:** Representative FACS plots of IFNγ and TNFα production in restimulated CD8 TILs from D12 MC38 tumors. Frequencies of cytokine producing CD8 TILs at indicated timepoints. Bars indicate median values. *P < 0.01, **P < 0.01, ***P < 0.001, ****P < 0.0001 by two-tailed unpaired *t* test (A, B, F, G), Mann-Whitney (C), Kruskal-Wallis test with Dunn’s correction (D), or one-way ANOVA with Tukey’s multiple comparisons test (E). Data is representative of two (B, C, D, G), three (E, F), or five (A) independent experiments.

We next determined whether the tumor resistance phenotype in *Lrp10^−/−^* mice depended on CD8 T cells. *Lrp10^+/+^*and *Lrp10^−/−^* were challenged with MC38 tumors and CD8 T cells were depleted *in vivo* through administration of an anti-CD8 antibody (Fig. 4D). Depleting CD8 T cells from *Lrp10^−/−^* mice eliminated their ability to resist the MC38 tumor. Together, these data show that *Lrp10^−/−^*mice accumulate higher levels of CD8 T cells within immunogenic tumors which are critical for enhanced tumor resistance.

### *Lrp10^−/−^* CD8 TILs maintain higher levels of IL7R and show reduced frequencies of terminally exhausted cells

IL7R expression on activated CD8 T cells marks cells with memory potential (Kaech *et al*, 2003) and is associated with enhanced responses to chronic viral infections and tumors (Belarif *et al*, 2018; Krishna *et al*, 2020; Micevic *et al*, 2023; Pauken *et al*, 2016). Given that Lrp10 downregulated IL7R during normal CD8 T cell homeostasis, we next compared IL7R levels on CD8 T cells in tumors from *Lrp10^+/+^* and *Lrp10^−/−^* mice to those from the spleen (Fig. 4E). Splenic CD8 T cells from *Lrp10^+/+^* and *Lrp10^−/−^* mice each displayed high levels of IL7R, with elevated levels observed in *Lrp10^−/−^*cells. Within the tumor, *Lrp10^+/+^* CD8 T cells showed >50% reduction in cell surface IL7R. In contrast there was no significant reduction in IL7R levels on *Lrp10^−/−^*CD8 TILs. Accordingly, tumors from *Lrp10^−/−^* mice showed greater frequencies of total IL7R+ CD8 TILs. These data demonstrate that loss of Lrp10 allows CD8 T cells to maintain higher levels of IL7R expression in the TME.

CD8 TILs are chronically exposed to antigen within the TME, which drives their differentiation into terminally exhausted cells (Chow *et al*, 2022; Giles *et al*., 2023). In a publicly available single cell RNAseq (scRNAseq) dataset of human CD8 T cells isolated from melanoma tumors (Sade-Feldman *et al*, 2018), Lrp10 gene expression overlapped with the expression of genes involved in T cell exhaustion and was negatively correlated with genes involved in stem and memory cell function (Fig. S5). Therefore, we next asked how *Lrp10* deletion affected the phenotype of CD8 TILs during the anti-tumor immune response.

We started by measuring expression of markers for memory cells (CD44 and CD62L) and for terminally exhausted cells (PD1 and Tim3) (Sakuishi *et al*, 2010; Zhou *et al*, 2011). *Lrp10^−/−^* mice accumulated increased frequencies of memory phenotype cells (CD62L+CD44+) in MC38 tumors both at early (day 12 after inoculation) and late (day 19 after inoculation) timepoints (Fig. 4F). While tumors in *Lrp10^+/+^* and *Lrp10^−/−^* mice harbored similar frequencies of terminally exhausted CD8 effectors (PD1+Tim3+) on day 12, *Lrp10^−/−^* mice showed reduced frequencies of PD1+Tim3+ CD8 effectors by day 19.

The ability to secrete inflammatory cytokines upon re-stimulation is a key characteristic of memory phenotype CD8 T cells. Therefore, we measured secretion of interferon-γ (IFN-γ) and tumor necrosis factor-α (TNF-α) in *Lrp10^+/+^* and *Lrp10^−/−^* CD8 TILs *in vitro* after re-stimulation with PMA/ionomycin (Fig. 4G). On day 12, the *Lrp10^−/−^* CD8 TIL population had ∼2.5-fold higher frequency of cells that secreted both IFN-γ and TNF-α compared to the corresponding *Lrp10^+/+^* population. Importantly, on day 19, the frequency of cytokine secreting *Lrp10^+/+^* CD8 T cells declined ∼3-fold while the *Lrp10^−/−^* population sustained levels that were similar to the earlier timepoint. Together, these data indicate that *Lrp10* deletion skews the composition of the CD8 TIL population away from terminal exhaustion toward a central memory phenotype that retains cytokine secretion capabilities.

### Single cell transcriptional and TCR profiling define CD8 TIL heterogeneity with and without Lrp10

We next sought to better define the identity and heterogeneity of *Lrp10^+/+^* and *Lrp10*^−/−^ CD8 TILs. *Lrp10^+/+^*and *Lrp10^−/−^* CD8 T cells were sorted from D12 MC38 tumors and subjected to paired scRNAseq and single cell TCR sequencing (scTCRseq). We analyzed quality transcriptome data from 1,269 *Lrp10^+/+^* cells and 6,267 *Lrp10^−/−^* cells. We performed unsupervised Louvain clustering of the pooled data, which yielded five distinct cell clusters visualized by uniform manifold approximation and projection (UMAP, Fig. 5A). Analysis of the most highly expressed genes between each cluster showed significant differences (Fig. 5B, Table S1). Cluster 0 showed high expression of naïve and memory genes (*Sell, CCr7, Tcf7, Lef1, Satb1, Bach2*). Cluster 1 showed high expression of genes associated with CD8 T cell effector function (*Gzma, Gzmb,* and *Ccl5*) and exhaustion (*Pdcd1 and Fasl*). Cluster 2 showed high expression of genes associated with innate CD8 T cells (*Klra7, Klrc2, Irak2, Fcer1g*). Cluster 3 showed high expression of genes associated with exhaustion (*Lag3, Rgs16, TNFRSF9*). Cluster 4 had very few cells of either genotype and showed high expression of the *Fos*, *Jun,* and *Id3* transcription factors.

**FIGURE 5:**
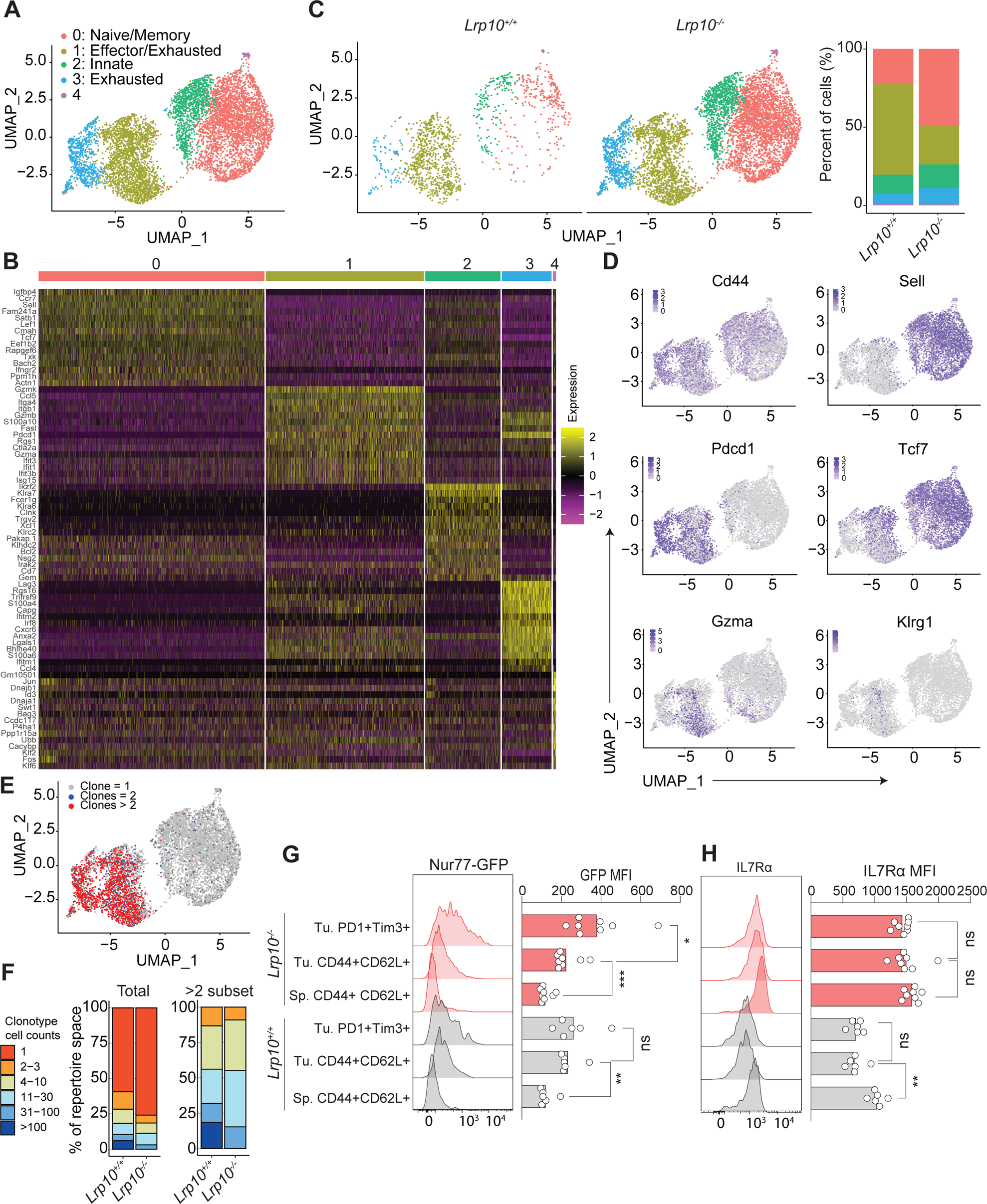
*Lrp10* deletion increases bystander CD8 T cell accumulation. **A:** UMAP of merged scRNAseq data from 1,269 *Lrp10^+/+^* cells and 6,267 *Lrp10^−/−^* CD8 TILs sorted from D12 MC38 tumors. Each dot corresponds to one individual cell. A total of 5 clusters (cluster 0 through 4) were identified and color-coded. **B:** A heatmap of the 15 most highly expressed genes in each cluster from panel A. Columns correspond to individual cells and rows correspond to genes. Color scale is derived from the z-score distribution from −2 (purple) to 2 (yellow). **C:** Contribution of *Lrp10^+/+^* and *Lrp10^−/−^* cells to the UMAP clusters identified in (A). **D:** Distribution of single cell transcript levels for *Cd44*, *Sell*, *Pdcd1*, *Tcf7*, *Gzma*, and *Klrg1* in the UMAP from panel A. Purple indicates gene expression and gray indicates no expression. **E:** Overlay of TCR clonotypes containing >2 cells, =2 cells, or =1 cell on the UMAP from (A). **F:** Stacked bar graph of relative clonotype abundance within the total CD8 TIL population and within the subset of clonotypes containing >2 cells. **G, H:** Representative FACS histograms of GFP and IL7Rα expression in splenic CD8 T_CM_ cells and CD8 TIL subpopulations from mice with D18 MC38 tumors. Bar graphs show GFP and IL7Rα MFI. Statistical significance was calculated using one-way ANOVA. Data represents two independent experiments.

While there was substantial overlap in the UMAP spaces occupied by *Lrp10^+/+^* and *Lrp10^−/−^* CD8 TILs, we noted key differences (Fig. 5C). *Lrp10^+/+^* cells fell predominantly into cluster 1 (exhausted/effector cells) while *Lrp10^−/−^*cells were found predominantly in cluster 0 (naïve/memory cells). Approximately equal frequencies of *Lrp10^+/+^* and *Lrp10^−/−^*cells were found in clusters 2, 3, and 4.

Cells in all clusters expressed *CD44*, indicating that they had at some point undergone activation (Fig. 5D). *Sell* expression was restricted to clusters 0 and 2, while *Pdcd1* expression was restricted to clusters 1 and 3. Expression of *Tcf7*, a central regulator of CD8 T cell memory differentiation and stem-like activity (Escobar *et al*, 2020; Pais Ferreira *et al*, 2020), was distributed between clusters 0, 1, and 2 and was largely excluded from cluster 3. Cells expressing the canonical effector genes *Gzma* and *Klrg1* were found primarily in cluster 1. scTCRseq showed that *Lrp10^+/+^*or *Lrp10^−/−^* CD8 T cells that had undergone clonal expansion (>2 cells detected per clonotype) were found almost exclusively in clusters 1 (exhausted/effectors) and 3 (exhausted) which closely overlapped with *Pdcd1* expression (Fig. 5E). In contrast, clusters 0 (naïve/memory cells) and 2 (innate-like cells) were predominantly composed of singlet clonotypes. The clonally expanded population in tumors from *Lrp10^+/+^*mice showed a few clonotypes that each contained a large number of cells. In contrast, the clonally expanded population in tumors from *Lrp10^−/−^* mice harbored a larger number of unique clonotypes that each contained relatively fewer numbers of cells (Fig. 5F). Interestingly, this finding is consistent with prior reports that enhanced IL7R signaling augments clonal diversity within responding CD8 T cell populations and reduces immunodominance (Melchionda *et al*, 2005; Sportes *et al*, 2008).

Together, transcriptional and TCR profiling show that *Lrp10* deletion enhances the accumulation of singlet CD8 T cells that express memory cell markers within the TME, reduces the overall frequency of cells expressing exhaustion markers, and reduces clonality within the clonally expanded TIL repertoire.

### All tumor infiltrating CD8 T cells show evidence of tumor reactivity

CD8 TILs are a heterogeneous population comprised of both tumor antigen-specific clones and bystander cells (Meier *et al*, 2022; Simoni *et al*, 2018). While bystander cells are associated with enhanced anti-tumor responses, the reasons for their accumulation and the mechanisms through which they act to influence tumor immunity are not well understood. The memory phenotype population that accumulated in tumors from *Lrp10^−/−^* mice did not show evidence of clonal expansion, therefore we speculated that they might be bystander cells.

To help identify tumor reactive CD8 T cells within MC38 tumors, we crossed *Lrp10^−/−^* mice to the *Nur77^GFP^* reporter strain, which exhibits GFP expression proportional to TCR stimulation (Au-Yeung *et al*, 2014; Moran *et al*, 2011). We challenged *Lrp10^+/+^;Nur77^GFP^* and *Lrp10^−/−^;Nur77^GFP^*mice with subcutaneous MC38 cells and harvested spleens and tumors on day 18 post-inoculation (Fig. 5G). GFP signal was generally absent in splenic CD8 T_CM_ cells and there was no Lrp10-dependent difference in GFP intensity. Compared to the splenic populations, GFP expression was increased in CD8 TILs from both *Lrp10^+/+^;Nur77^GFP^* and *Lrp10^−/−^;Nur77^GFP^* mice, indicating heightened tumor reactivity. GFP signal from CD8+CD62L+CD44+ and CD8+PD1+Tim3+ cells were similar in tumors from *Lrp10^+/+^;Nur77^GFP^* mice. In contrast, CD8+PD1+Tim3+ cells from the *Lrp10^−/−^;Nur77^GFP^*TIL population displayed increased levels of GFP signal compared to CD8+CD62L+CD44+ cells, suggesting that high levels of TCR signaling were required to drive *Lrp10^−/−^* CD8 T cells to a terminally differentiated phenotype.

Together, these data show that the majority of CD8 T cells infiltrating tumors have some degree of tumor reactivity regardless of Lrp10 status. Significantly, despite not undergoing clonal expansion, central memory phenotype CD8 TILs from *Lrp10^−/−^* mice showed evidence of active TCR signaling and thus were not tumor-ignorant, inactive bystanders.

TCR signaling downregulates IL7R expression (Chandele *et al*, 2008). Accordingly, GFP+ CD8 TILs from *Lrp10^+/+^;Nur77^eGFP^*mice showed a decline in cell surface expression of IL7R compared to splenic T_CM_ cells. Notably, CD8 TILs from *Lrp10^−/−^;Nur77^eGFP^*mice showed sustained IL7R expression, even in terminally differentiated PD1+Tim3+ cells, suggesting that *Lrp10* deletion uncouples IL7R expression from chronic TCR signaling (Fig. 5H).

### *Lrp10* deletion skews the clonally expanded population away from exhaustion

We next assessed how *Lrp10* deletion affected the differentiation of clonally expanded CD8 T cells in MC38 tumors. We analyzed differential gene expression between *Lrp10^+/+^* and *Lrp10^−/−^* CD8 TILs for clonotypes that contained more than two cells, corresponding to a subpopulation of cells within clusters 1 and 3 in the original UMAP (Fig. 6A, Table S2). Among genes upregulated in clonally expanded *Lrp10^−/−^* CD8 TILs were those commonly associated with interferon responses (*Ifitm1*, *Ifitm2*, *Ifitm3*, *Plac8*, *Capg*) and cytotoxicity (*Jaml*, *Ctsw*, *Klrk1*). In contrast, *Lrp10^+/+^* clonally expanded cells showed increased expression of epigenetic regulators associated with T cell exhaustion (*Tox*) and hematopoietic differentiation (*Jarid2*) (Kinkel *et al*, 2015) as well as *Fgfr2* and *Hexb*. We did not note any difference in the expression of stem-memory genes (*IL7R, Tcf7*) or in other transcriptional regulators of exhaustion (*Tbx21, Eomes*, *Prdm1*) (Fig. 6B).

**FIGURE 6:**
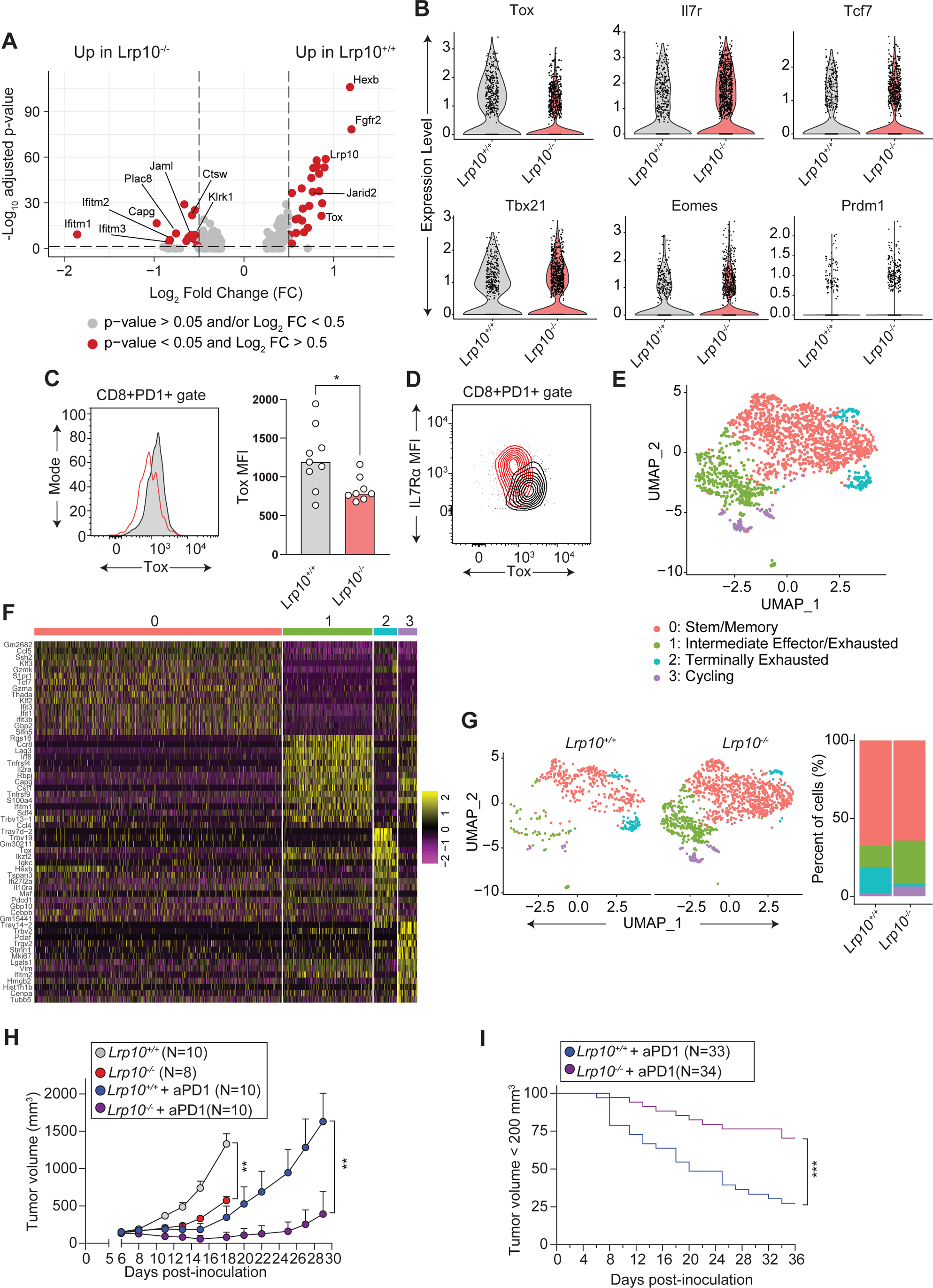
*Lrp10* deletion counteracts exhaustion programming in clonally expanded CD8 TILs. **A:** Volcano plot showing differentially expressed genes in *Lrp10^+/+^* and *Lrp10^−/−^* CD8 TIL clonotypes containing > 2 cells. **B:** Violin plots of *IL7R*, *Tox*, *Tcf7*, *Fgfr2*, *Tbx21, Eomes, Prdm1*. **C:** Representative FACS histogram of Tox expression and quantification of Tox expression in D12 CD8+PD1+ TILs. Data is combined from two independent experiments. **D:** Representative FACS plot of Tox vs IL7R expression in CD8+PD1+ TILs from the mice in (C). **E:** UMAP of reclustered scRNAseq data from 1,673 CD8 TILs containing clonotypes with more than 2 cells. Each dot corresponds to one individual cell. A total of 4 clusters (cluster 0 through 3) were identified and color-coded. **F:** A heatmap of the 15 most highly expressed genes in each cluster from panel E. Columns correspond to individual cells and rows correspond to genes. Color scale is derived from the z-score distribution from −2 (purple) to 2 (yellow). **G:** Contribution of *Lrp10^+/+^* and *Lrp10^−/−^* cells to the UMAP clusters identified in (E). **H:** MC38 tumor growth in *Lrp10^+/+^* and *Lrp10^−/−^*mice with and without anti-PD1. Data presented as mean with SEM at each point. Statistical differences between treatment groups were calculated using a Mann-Whitney test at the indicated time points. Results are representative of three independent experiments. **I:** Frequency of tumor progression in *Lrp10^+/+^* and *Lrp10^−/−^*mice treated with anti-PD1 across three separate cohorts. Mice were said to have progressed for volumes >200 mm^3^. *P < 0.01, **P < 0.01, ***P < 0.001, ****P < 0.0001 by two-tailed unpaired *t* test (C), Mann-Whitney (H), and log-rank (Mantel-Cox) test (I).

Given the importance of *Tox* in CD8 T cell exhaustion (Khan *et al*, 2019; Scott *et al*, 2019; Seo *et al*, 2019; Yao *et al*, 2019), and its apparent upregulation in clonally expanded *Lrp10^+/+^*CD8 TILs, we specifically measured Tox protein expression in CD8 TILs in day 12 MC38 tumors (Fig. 6C, D). We used PD1 expression as a surrogate marker for clonally expanded CD8 TILs given the high degree of overlap of between *Pdcd1* expression and clonotypes containing two or more cells. Within the PD1+ subpopulation, *Lrp10^−/−^*cells CD8 TILs displayed lower levels of *Tox* expression, consistent with our scRNAseq analysis. Comparing IL7R and Tox levels between PD1+ *Lrp10^+/+^*and *Lrp10^−/−^* CD8 TILs showed that the *Lrp10^−/−^*population was skewed toward increased IL7R expression and decreased Tox expression whereas this relationship was inverted in *Lrp10^+/+^* cells. These data support the conclusion that *Lrp10* deletion attenuates CD8 TIL terminal exhaustion in tumors and suggests that Lrp10 potentiates Tox-induced exhaustion programming.

Reclustering the subset of clonotypes containing 2 or more cells resulted in a UMAP partitioned into four subclusters (Fig. 6E). Analysis of the differentially expressed genes between each cluster showed high expression of *Tcf7* in cluster 0, together with effector genes like *Ccl5* and *Gzma* (Fig. 6F, Table S3). This profile suggests that cluster 0 represents stem/memory anti-tumor CD8 T cells. Cluster 1 showed high expression of genes associated with exhaustion (*Lag3*, *Rgs16*, *Tnfrsf9*) and effector function (*Irf8*, *Tnfrsf4*, *Il2ra*, *Ccl4*), indicating an intermediate effector/exhausted phenotype. In contrast, cluster 2 showed upregulation of multiple factors known to facilitate terminal exhaustion (*Tox*, *Ikzf2, Maf*). Cluster 3 showed upregulation of cell cycle genes (*MKi67*, *Pclaf, Stmn1*, *Tubb5, Hmgb2*) indicating a sub-population of proliferative cells.

Most *Lrp10^+/+^* and *Lrp10^−/−^* cells were in cluster 0 (stem/memory) where they occurred at approximately equal frequencies (Fig. 6G). *Lrp10^−/−^* cells also partitioned into clusters 1 (intermediate effectors/exhausted) and 3 (cycling). In contrast, after cluster 0, most *Lrp10^+/+^* cells were found in cluster 2 (terminally exhausted). *Lrp10^+/+^* cells were found to a lesser extent in cluster 1 and were absent from cluster 3. Together, these data suggest that *Lrp10* deletion reduces the frequency of terminal exhaustion amongst clonally expanded CD8 TILs and, instead, enriches cells with an intermediate effector/exhausted phenotype or a proliferative phenotype.

### *Lrp10* deletion synergizes with anti-PD1 immunotherapy to cure tumors at high frequencies

The frequency of CD8 terminal exhaustion in the TME is inversely correlated with the efficacy of immune checkpoint inhibition in chronic viral infections and cancer (Kurtulus *et al*, 2019; McLane *et al*, 2019; Miller *et al*, 2019; Sade-Feldman *et al*., 2018). The immunological and transcriptional phenotype of *Lrp10^−/−^* CD8 TILs suggested that they were resistant to terminal exhaustion and instead primarily expressed effector or memory features. Therefore, we hypothesized that *Lrp10* deletion would enhance the effect of immune checkpoint inhibition. We injected *Lrp10^+/+^* and *Lrp10^−/−^*mice with 2.5 mg/kg of anti-PD1 for three doses starting on day 6 after inoculation with MC38 cells (Fig. 6H). Anti-PD1 at this dose imparted partial MC38 resistance to *Lrp10^+/+^* mice which resembled the tumor growth rate in *Lrp10^−/−^* mice treated with vehicle. In contrast, anti-PD1 treatment strongly synergized with *Lrp10* deletion to slow the growth of MC38 tumors. Over several cohorts of mice, we found that anti-PD1 treatment coupled with *Lrp10* deletion resulted in tumor rejection more frequently compared to anti-PD1 treatment alone (Fig. 6I). Thus, *Lrp10* deletion imparts both partial intrinsic tumor resistance and enhanced susceptibility to immune checkpoint inhibition.

## DISCUSSION

Through forward genetic screening in mice, we discovered that *Lrp10* maintains CD8 T cell homeostasis by limiting the size of the naïve and central memory subpopulations. Our data suggest a model where Lrp10 downregulates IL7R to curtail the number of CD8 T cells that survive in peripheral lymphoid organs. We propose that induction of Lrp10 during CD8 T cell stimulation reduces the number of cells able to compete for IL7 and enter the central memory repertoire. We hypothesize that this mechanism prevents the propagation of long-lived memory cells expressing potentially self-reactive antigen receptors. Consequently, Lrp10 may help drive CD8 T cell responses toward immunodominant antigens by suppressing the number and diversity of cells that persist after clonal contraction. Notably, *Lrp10^−/−^* mice are not overtly autoimmune. *Lrp10^−/−^* mice are currently housed under specific pathogen free conditions in which the environmental antigen burden is low (Beura *et al*, 2016). Whether exposing *Lrp10^−/−^* mice to “dirtier” environments would amplify central memory populations with the potential to cause autoimmune tissue damage remains to be tested. Furthermore, the factors that control Lrp10 expression are incompletely understood. Defining Lrp10’s expression and functional responses to TCR signal strength, co-stimulation, and inflammatory cytokines would provide insights into how Lrp10 integrates diverse external signals to shape CD8 T cell fate and the composition of the central memory repertoire.

Deletion of *Lrp10* also imparts intrinsic resistance to immunogenic tumors. This phenotype depends on extensive infiltration of CD8 T cell populations that harbor reduced frequencies of terminally exhausted cells, retain high levels of IL7R and other memory cell characteristics, and display enhanced susceptibility to immune checkpoint inhibition. Our data suggest two ways in which *Lrp10* deficiency influences anti-tumor immunity. First, it impairs terminal exhaustion of clonally expanded anti-tumor CD8 T cells. We showed that lack of Lrp10 suppresses the frequency cells expressing *Tox*. The mechanism that underpins this finding remains unclear. We hypothesize that elevated IL7R expression on *Lrp10^−/−^* CD8 T cells helps them resist terminal differentiation. One possibility is that enhanced IL7R signaling directly counteracts Tox, for example through activated STAT5 signaling networks (Ding *et al*, 2020). In this scenario, rational combination of IL7R agonists with immune checkpoint inhibition may benefit immunotherapy approaches. Our data also suggest that Lrp10 acts as a brake on IL7R signaling and thus may limit IL7-based therapies. Whether the ability of *Lrp10* deletion to impair T cell exhaustion depends exclusively on IL7R, or whether other signaling networks are involved, remains to be tested.

A second way *Lrp10* deletion may promote anti-tumor immunity is through the accumulation of numerous singlet CD8 clones with a memory phenotype within the TME. Based on their lack of clonal amplification and absent PD1 expression, these cells appeared to be tumor-ignorant bystanders. However, they show TCR signaling above background indicating that they are enriched for tumor reactivity. One possibility is that these CD8 T cells express TCRs that are cross-reactive with tumor antigens and endogenous or microbial antigens, a population called “false bystanders” (Bessell *et al*, 2020; Chiou *et al*, 2021; Meier *et al*., 2022). *Lrp10^−/−^* mice may inherently possess high frequencies of these cells due to upregulated IL7R permitting CD8 T cells with cross-reactive TCRs to persist in the central memory pool. Cross-reactive CD8 T cells are frequently found in human tumors and mouse cancer models (Caushi *et al*, 2021; Danahy *et al*, 2020; Simoni *et al*., 2018). There is clinical interest in exploiting these cells for functional anti-tumor responses (Batich *et al*, 2017; Millar *et al*, 2020; Rosato *et al*, 2019). Our data suggest Lrp10 is part of a previously unrecognized genetic program that limits the number of false bystanders in tumors to suppress intrinsic anti-tumor immunity.

We have shown that Lrp10 suppresses IL7R at a post-translational level. Forced expression of Lrp10 impairs IL7R glycosylation and maturation through protein-protein interactions. Lrp10 is an orphan endocytic receptor with no known ligands. Thus, exactly how Lrp10 limits cell surface IL7R in primary CD8 T cells remains unclear. It will be important to define the intracellular trafficking itinerary and subcellular localization of Lrp10 relative to IL7R as a function of cell differentiation state. Additionally, the discovery of ligands, or co-receptors, that control Lrp10 function could reveal specific environmental cues that guide the differentiation and fate of activated T cells through IL7R signaling.

The number of NK cells was reduced in *Lrp10^−/−^* mice with a corresponding deficit in *in vivo* NK cytolytic activity. The reason for this deficiency is currently unknown. T cells and NK cells compete for limiting amounts of IL7 for survival (Carrette & Surh, 2012; Martin *et al*, 2017). It is possible that increased IL7R expression on T cells deprives NK cells of adequate IL7 to ensure their survival. Alternatively, Lrp10 may have a cell-intrinsic role in the development or maintenance of the NK cell lineage. Specific deletion of Lrp10 in T cells versus NK cells would help define the role for Lrp10 in NK cell homeostasis.

Through adoptive transfer studies, we demonstrated that Lrp10 acts in a cell-autonomous manner to limit IL7R expression and suppress CD8 T cell homeostatic expansion. One significant limitation of the current study is the use of a constitutive knockout model to define Lrp10’s function in anti-tumor immune responses. While this approach can approximate what happens during global inhibition of Lrp10 in a therapeutic setting, it does not rule out that loss of Lrp10 in other cell populations (e.g., stromal, myeloid, or CD4 T cells) may influence CD8 T cell function. Additionally, although *Lrp10* deletion did not distort thymic cellularity, it may subtly change thymic T cell developmental trajectories or TCR selection criteria to render mice tumor resistant. Specifying the lineages and timeframe in which *Lrp10* is deleted would help further define its role in anti-tumor immunity. Another limitation of this study is our finding that the tumor resistance phenotype of *Lrp10^−/−^* mice was restricted to the highly immunogenic MC38 tumor. Currently, it is not clear what tumor factors dictate immune responsiveness in the setting of *Lrp10* deletion. We hypothesize that a high tumor mutational burden is important for generating the increased polyclonality observed amongst *Lrp10^−/−^*CD8 TILs. Future studies will define how tumor mutational burden affects anti-tumor immunity, and the clonality and differentiation phenotypes of CD8 TILs, in the context of *Lrp10* deletion.

In summary, we present Lrp10 as a new determinant of IL7R expression that has important implications for CD8 T cell fate decisions during normal homeostasis and anti-tumor immune responses.

## MATERIALS AND METHODS

### Mouse strains

Mice were housed in specific pathogen-free conditions and fed a normal chow diet at the University of Texas Southwestern Medical Center. All animal experiments were performed according to institutionally approved protocols. ENU-mutagenesis, strategic breeding of mutagenized mice, phenotypic screening, and automated meiotic mapping were performed as previously described. *B6 CD45.1*, *Nur77^GFP^*, *OT1*, and *B2M^−/−^*strains were obtained from the Jackson Laboratory. These strains were intercrossed with *Lrp10^−/−^* mice as needed. Male and female mice aged 8-16 weeks were used for experiments.

### Generation of knockout mouse strains using the CRISPR/Cas9 system

To generate single knockout mouse strains, female C57BL/6J mice were super-ovulated by injection of 6.5 U pregnant mare serum gonadotropin (PMSG; Millipore), followed by injection of 6.5 U human chorionic gonadotropin (hCG; Sigma-Aldrich) 48 h later. The super-ovulated mice were subsequently mated overnight with C57BL/6J male mice. The following day, fertilized eggs were collected from the oviducts. *In vitro*-transcribed Cas9 mRNA (50 ng/µl) and *Lrp10* small base-pairing guide RNA (50 ng/µl) were injected into the cytoplasm or pronucleus of embryos. Injected embryos were cultured in M16 medium (Sigma-Aldrich) at 37°C in 5% CO2. For the production of mutant mice, two-cell stage embryos were transferred into the ampulla of the oviduct (10–20 embryos per oviduct) of pseudo-pregnant Hsd:ICR (CD-1) female mice (Harlan Laboratories). An *Lrp10* CRISPR allele that resulted in a frame-shifting 8 bp deletion in exon 5 was used for all experiments. *Lrp10* deletion was verified at the protein level through Western blotting.

### Plasmids

Full length mouse Lrp10 and Lrp10-ECD were tagged with a C-terminal HA epitope for use in co-immunoprecipitation experiments. For retroviral complementation experiments, full length Lrp10 was sub-cloned into MSCV-IRES-GFP. Mouse CD127 with a C-terminal FLAG tag was obtained from SinoBiological. Details of plasmids are available upon request. MSCV-IRES-GFP was a gift from Tannishtha Reya, Addgene plasmid # 20672.

### Adoptive transfer experiments and *in vivo* CD8 T cell functional assays

For homeostatic proliferation experiments, recipient mice were sub-lethally irradiated with 6 Gy given as a single dose 24h prior to cell transfer (X-RAD 320, Precision X-ray). Naïve CD8 T cells were purified to >90% from the spleens of donor strains using negative selection (StemCell Technologies) and stained with CTV or CTFR proliferation dyes (Molecular Probes) according to the manufacturer’s instructions. Labelled cells were combined at a 1:1 ratio and 1-2 million cells were injected intravenously into recipients. For IL7R blockade, recipient mice were given intraperitoneal injections of 500 µg anti-IL7R (clone A7R34, BioXCell) or isotype control (clone 2A3, BioXCell) on days 1, 3, and 5 post-cell transfer. In immunization experiments, unirradiated mice were given a single intraperitoneal injection of 100 µg SIINFEKL (Vivitide) or SIITFEKL (ANASPEC) 24h after cell transfer. Spleens were harvested at the indicated time points for analysis by flow cytometry. The frequency and proliferation status of donor cells were measured based on the indicated markers and the dilution of the proliferation dyes. *In vivo* CD8 and NK cytotoxicity assays were performed as previously described (Choi *et al*, 2019).

### Tumor models

C57Bl/6 syngeneic MC38 (ATCC) and B16F10 (ATCC) cells were maintained in DMEM with 10% FBS and passaged three times per week. Cells were discarded after 30 passages. Mice 8-16 weeks of age were injected subcutaneously with 5 x 10^5^ (MC38) or 4 x 10^5^ (B16F10) cells on the right flank. Approximately equal numbers of male and female mice of each genotype were used in experiments. Tumor volume was measured three times weekly starting on day 6 post-inoculation and was calculated based on the formula *length* x *width* x *width/2*. For immune checkpoint inhibition experiments, mice were given intraperitoneal injections of 2.5 mg/kg anti-PD1 (clone RMP1-14, BioXCell) on days 5, 8, and 11 post-tumor inoculation. For CD8 depletion experiments, mice were given intraperitoneal injections of 10 mg/kg anti-CD8 (clone YTS 169.4, BioXCell).

### CD8 TIL isolation and flow cytometry

At the indicated time points, tumors were harvested, minced with razor blades in RPMI, and digested using the Miltenyi Tumor Dissociation Kit for 1h at 37°C according to manufacturer’s protocol, except for reducing the amount of enzyme R by 90% to preserve cell surface epitopes. Dissociated tumors were passed through 70 µm filters and the volume of tumor cell suspension equivalent to 150 µg of tumor was stained with cell viability dye and the indicated cell surface markers. For cytokine production analyses, the volume of tumor cell suspension equivalent to 150 µg of tumor was stimulated with a cocktail of PMA, ionomycin, and Brefeldin A for 4h at 37°C followed by staining live cell dye and antibodies against cell surface markers. Cells were then fixed and permeabilized with the Intracellular Fixation & Permeabilization Buffer Set (eBioscience) and then stained with fluorochrome-conjugated anti-cytokine antibodies. The Mouse FoxP3 Fixation and Permeabilization Buffer Set (BDBiosciences) was used for Tox intracellular staining. Flow cytometry data was collected on an LSR Fortessa (BDBiosciences) and analyzed using FlowJo software.

### Transfection, co-immunoprecipitation assays, and Western blotting

HEK293T cells were maintained in DMEM containing 10% FBS and routinely tested for mycoplasma (Fisher Scientific). Cells were transfected in 6-well plates with 2 µg of the indicated constructs in the presence of Lipofectamine 2000 (Invitrogen) according to the manufacturer’s instructions. At 48 h post-transfection, cells were rinsed in cold PBS and lysed in buffer containing 1% NP-40 and HALT protease inhibitor (Thermo) followed by centrifugation at 13,000 rcf for 10 min. Co-IP of FLAG-tagged proteins was performed by incubating M2 anti-FLAG resin (Sigma) with clarified cell lysates for 2 h at 4°C with end-over-end rotation. Beads were washed four times in cold lysis buffer, and protein complexes were eluted with 150 mg/ml of 3X FLAG peptide (Sigma). Samples were diluted in 2X SDS sample buffer and analyzed with SDS–PAGE according to standard procedures. For analysis of differential IL7Ra glycosylation, transfected cells were lysed, sonicated, and boiled in 200 µL of buffer containing 1% SDS, HALT protease inhibitor, and benzonase (Sigma). Denatured lysates were then diluted to 1.5 mL in 1% NP-40 and anti-FLAG IP was performed as usual. Eluted proteins were then treated with PNGase F (Promega) according to manufacturer’s protocol and analyzed with SDS-PAGE. For Western blotting on primary cells, cell pellets were lysed in buffer containing 1% SDS, HALT protease inhibitor, and benzonase. Protein levels were normalized using the bicinchoninic acid (BCA) assay (Pierce) and 10–15 µg of protein was diluted in 4X SDS sample buffer and analyzed with SDS– PAGE.

### Real-time quantitative PCR measurements

RNA from purified splenic CD8 T cells was reverse transcribed into cDNA using oligo-d(T) primers and M-MuLV reverse transcription (Promega). Real-time quantitative PCR was performed using SYBR DNA polymerase (Thermo Scientific) and target specific primers.

### scRNAseq and scTCRseq

1.5 x 10^4^ live CD8 T cells were FACS sorted from dissociated D12 MC38 tumors. scRNAseq and scTCRseq libraries were generated using the Chromium Next GEM Single Cell 5’ kit (10X Genomics) according to the manufacturer’s protocol. Purified libraries sequenced together on one S4 lane of a NovaSeq sequencing instrument with the run configuration 150×10×10×150.

### scRNAseq analysis

scRNAseq FASTQ files were processed with Cell Ranger v. 7.1.0 (10x Genomics) using default settings for 5’ RNA gene expression analysis. Using Seurat (v. 4.4.0) in R, we selected for high quality transcriptomes by filtering with the following criteria: 500 to 6000 features (detected genes), 2,000 to 40,000 unique modular identifier (UMI) counts, ribosomal gene content between 5 and 50%, and mitochondrial gene content below 5%. Next, we used the scGate package to select target cells that were Cd8a and Cd8b1 positive, resulting in 1372 and 6775 high quality *Lrp10^+/+^* and *Lrp10^−/−^* TIL transcriptomes, respectively. 103 *Lrp10^+/+^* and 508 *Lrp10^−/−^* doublets were predicted and removed using DoubletFinder. Next, we pooled the data (7536 transcriptomes) from the filtered feature matrices and added the corresponding scTCRseq clonotype data into the metadata of the Seurat objects using the filtered contig annotations and clonotype csv files. After removal of a cluster of contaminating cells expressing NK markers, we performed standard normalization and scaling of the remaining 7333 transcriptomes and identification of 1000 highly variable genes (HVGs) using the vst method in Seurat. Dimensionality reduction on the HVGs was achieved using principal component analysis (PCA) and UMAP on the first 15 principal components. Unsupervised clustering using the Louvain algorithm was implemented using the FindNeighbors and FindClusters functions in Seurat with resolution 0.2. Differentially expressed genes (DEGs) between clusters were identified using FindAllMarkers with the parameters min.pct = 0.25, logfc.threshold = 0.25 and excluding mitochondrial, ribosomal, heat shock, and cell cycle genes. Using the dplyr package and DoHeatmap, we generated a heatmap of the top 15 genes by average fold change in each cluster. Distributions of individual genes signatures and clonotypes in the UMAP were visualized using the FeaturePlot function. Stacked bar graphs were generated using the dittoBarPlot function in the dittoSeq package (https://rdrr.io/bioc/dittoSeq/). The subset of 1,673 transcriptomes with clonotypes greater than 2 was extracted and clustered using the first 30 principal components at 0.25 resolution. FindMarkers was used to find the DEGs between the *Lrp10^+/+^* and *Lrp10^−/−^* cells and FindAllMarkers was used to identify DEGs between clusters. We used EnhancedVolcano (https://github.com/kevinblighe/EnhancedVolcano) with fold change cutoff of 0.5 and p value cutoff of 0.05 to visualize the DEGs between the *Lrp10^+/+^* and *Lrp10^−/−^* cells.

### scTCRseq analysis

scTCRseq FASTQ files were processed with Cell Ranger v. 7.1.0 (10x Genomics) using default settings with the Single Cell V(D)J-T (Alpha Beta) library. We analyzed clonal proportions using the repClonality function with the “rare” method in the Immunarch 1.0.0 R package (ImmunoMind) to segregate the data according to default bin settings (clonotype counts = 1, 2-3, 4-10, 11-30, 31-100, 101-MAX).

### Generation of anti-Lrp10 monoclonal antibodies

To generate anti-Lrp10 monoclonal antibodies, *Lrp10^−/−^*mice were immunized with recombinant murine Lrp10-ECD in alum adjuvant followed by two protein-only boosts every two weeks. Anti-Lrp10 serum titers were monitored by ELISA. After the final boost, the spleen of the mouse with the highest antibody titers was dissociated and fused with Sp2/0-Ag14 myeloma cells (ATCC) using the ClonaCell-HY Hybridoma Kit (StemCell Technologies) according to the manufacturer’s protocol. The fusion product was plated in semi-solid selection media and clonal outgrowths were harvested into single wells of 96 well plates after 10-14 days. Single clones were allowed to grow for another 14 days in suspension culture after which the supernatant was tested for reactivity against Lrp10-ECD using ELISA. Wells that scored positive in the ELISA assay were then evaluated by flow cytometry against cell lines that overexpressed full-length Lrp10. Wells that scored positive in the ELISA and FACS assays were propagated and rescreened. Three different hybridomas that produced anti-Lrp10 antibodies were generated from a single mouse of which only one (clone 6H5) could detect the protein by Western blot. Monoclonality of this hybridoma was verified by sequencing. Antibody was purified using a Hi-Trap Protein G column (Cytiva). Notably, although all three antibodies were screened for FACS reactivity on intact cells, they could only detect cell surface Lrp10 when it was overexpressed. None of them could detect cell surface Lrp10 on primary CD8 T cells.

### Statistical analysis

No statistical methods were used to pre-determine sample size. Normal distribution of data was determined by the Shapiro–Wilk normality test. For normally distributed data, the statistical significance of differences between experimental groups was determined by paired or unpaired t tests as indicated. For non-normally distributed data, a non-parametric test was used as indicated. Multiple comparisons were analyzed with ANOVA. Statistical analyses were performed using GraphPad Prism software. Differences with P values < 0.05 were considered significant. P values are denoted by *P < 0.05, **P < 0.01, and ***P < 0.001. Differences with P values ≥ 0.05 were considered not significant (ns).

## ACKNOWLEDGEMENTS

We thank Emre Turer, Anne Satterthwaite, and Tuoqi Wu for critical reading of this manuscript. We thank Sook Kyung Chang and Elsam Elghonaimy for helpful discussions. We also thank Kennith Stedham for his help with managing the mouse colony. We acknowledge Caitlin Eaton and Vanessa Schmid from the UTSW McDermott Next Generation Sequencing Core for library preparation and sequencing during the single cell experiments. Initial portions of this work were funded by a sponsored research agreement with ImmunoDesigners, Inc. This work was funded by NIH R01-AI125581 (B.B.) and NIH R01-AI167920 (E.N.-G.).

## AUTHOR CONTRIBUTIONS

Conceptualization: E.N.-G. and B.B.; Methodology: L.C., A.L., J.W., B.B., and E.N.-G.; Experiments: J.R., A.L., J.W, S.G., X.Z., H.S., and E.N.-G.; Data analysis: L.C. and E.N.-G.; Visualization: L.C. and E.N.-G.; Writing: E.N.-G.; Funding acquisition: B.B. and E.N-G.; Supervision: E.N.-G.

---

**FIGURE S1:**
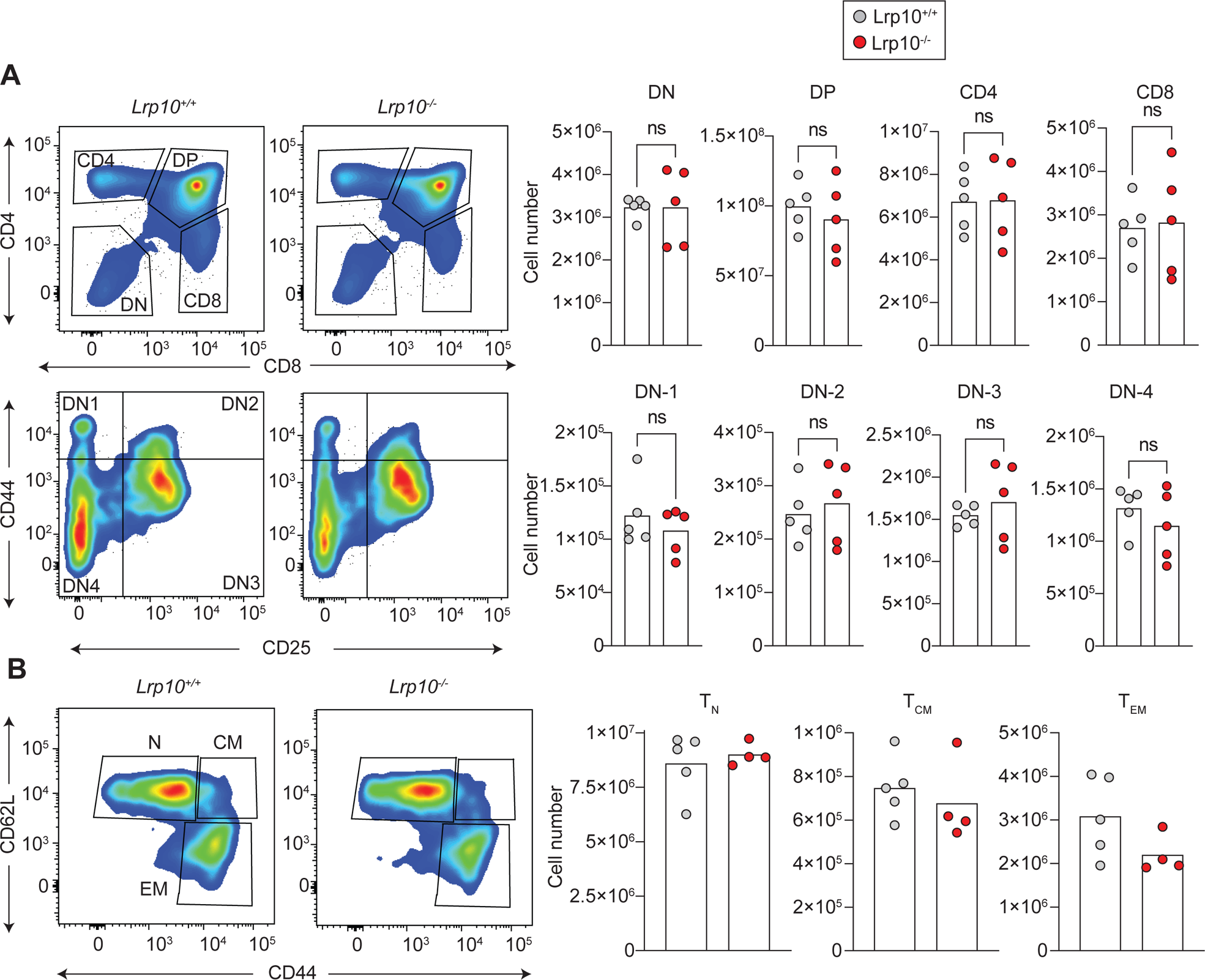
**A:** Representative FACS plots and numbers of thymic T cell subpopulations. **B:** Representative FACS plots and numbers of splenic CD4 subpopulations.

**FIGURE S2:**
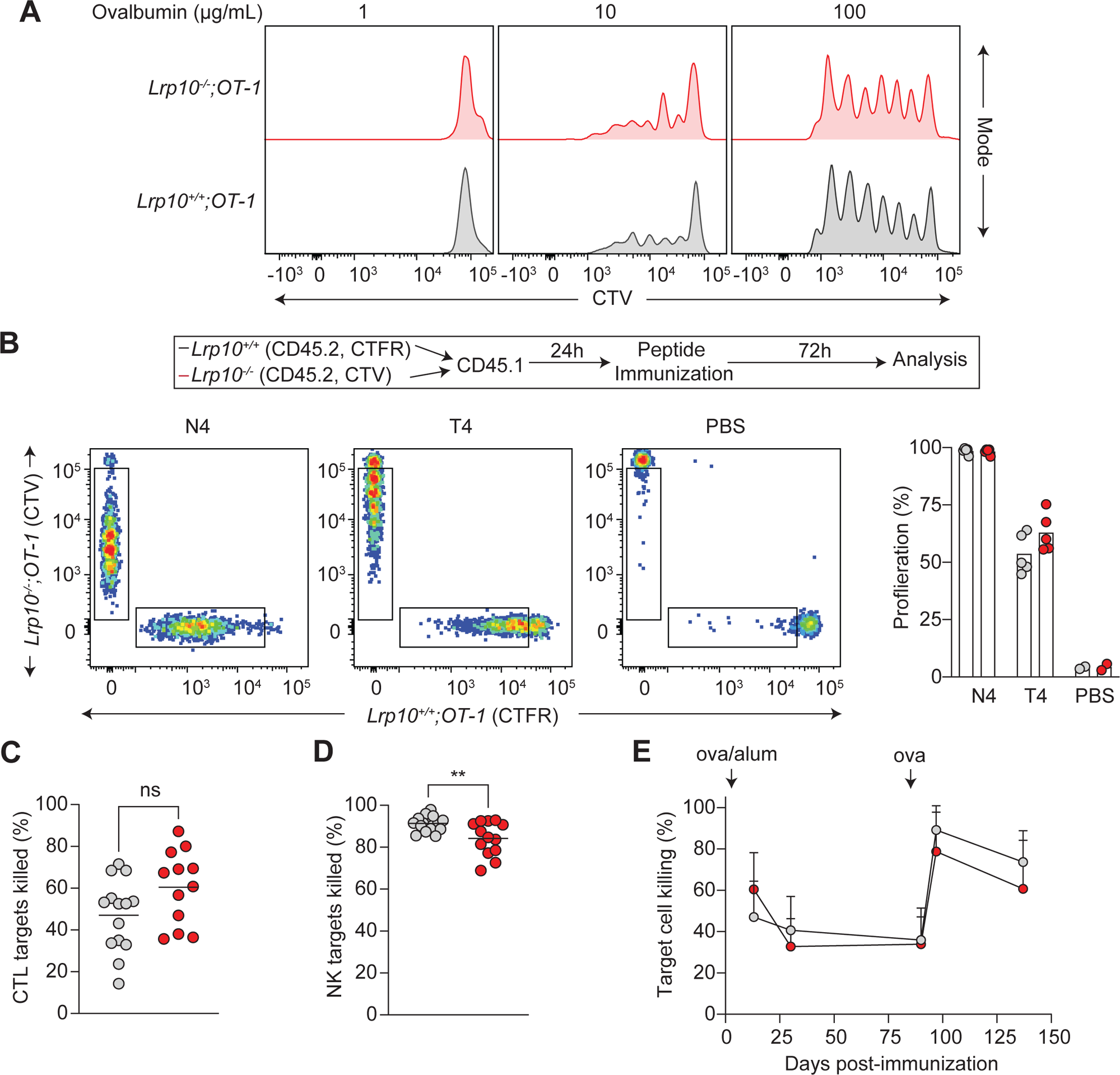
**A:** Proliferation of *Lrp10^+/+^;OT-1* or *Lrp10^−/−^;OT-1* cells 72h after co-culture with *Lrp10^+/+^* dendritic cells pulsed with the indicated concentrations of ovalbumin. **B:** 10^6^ *Lrp10^+/+^;OT-1* and *Lrp10^−/−^;OT-1* cells were labeled with CTFR and CTV, respectively, and injected into unirradiated recipients. 24h later mice were immunized with SIINFEKL (N4), SIITFEKL (T4), or PBS. Cell proliferation was measured 72h after immunization based on dye dilution. **C:** *In vivo* CTL cytotoxicity in *Lrp10^+/+^* and *Lrp10^−/−^*mice 12 days after immunization with ovalbumin and aluminum adjuvant (ova/alum). **D:** *In vivo* NK cytotoxicity assay in *Lrp10^+/+^* and *Lrp10^−/−^* mice injected with labeled MHC-I deficient target cells. **E:** Serial *in vivo* cytotoxicity assays in the mice from (E) after immunization with ova/alum. Mice were given a boost of ova protein alone on day 90.

**FIGURE S3:**
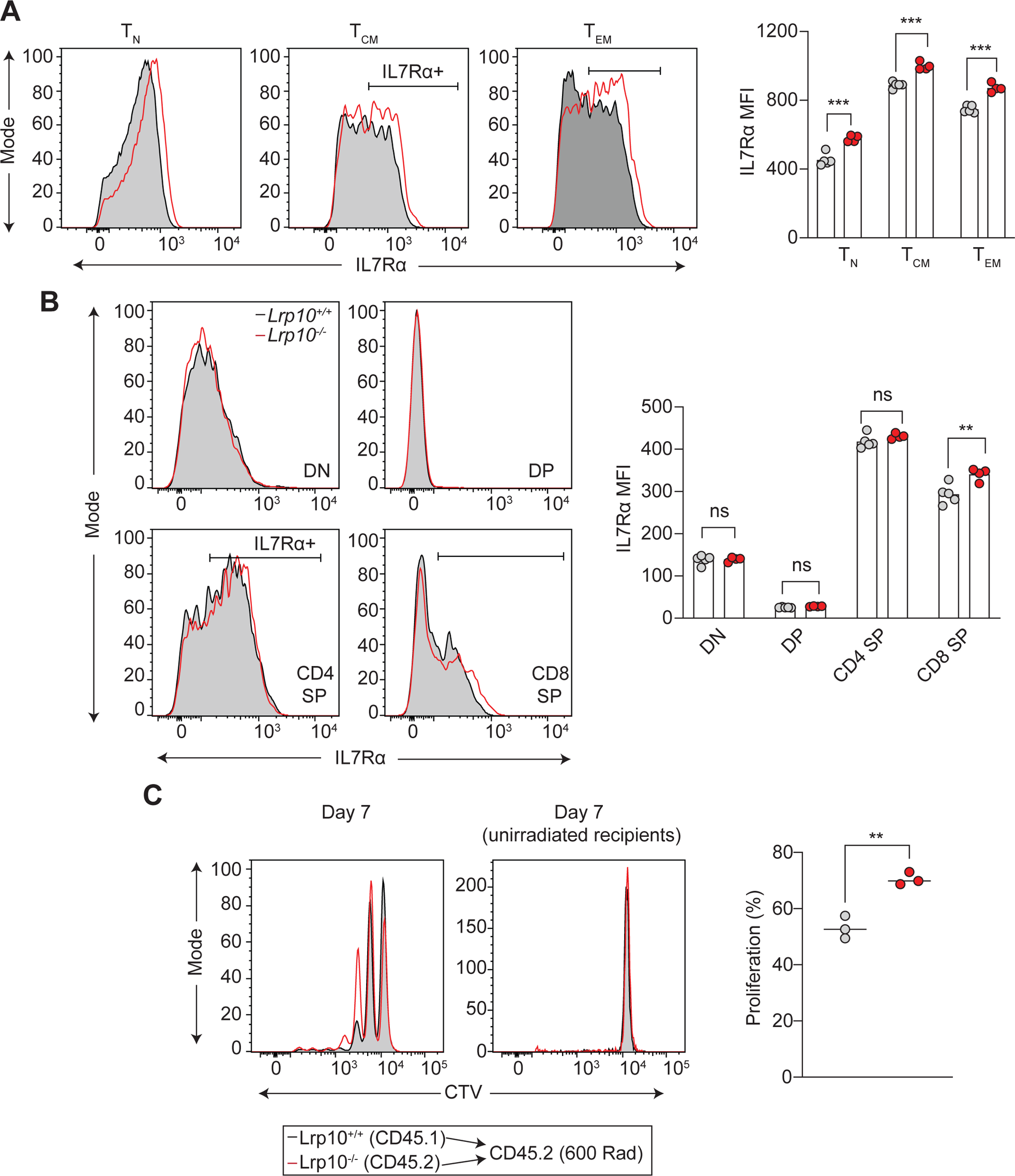
**A:** Cell surface IL7R expression in CD4 subpopulations. **B:** Cell surface IL7R expression on thymic T cell subpopulations. **D:** Homeostatic expansion of *Lrp10^+/+^* and *Lrp10^−/−^*CD4 T cells labeled with CTV and injected in sub-lethally irradiated and unirradiated recipients. Proliferation was measured based on the fraction of cells that underwent at least one cell division.

**FIGURE S4:**
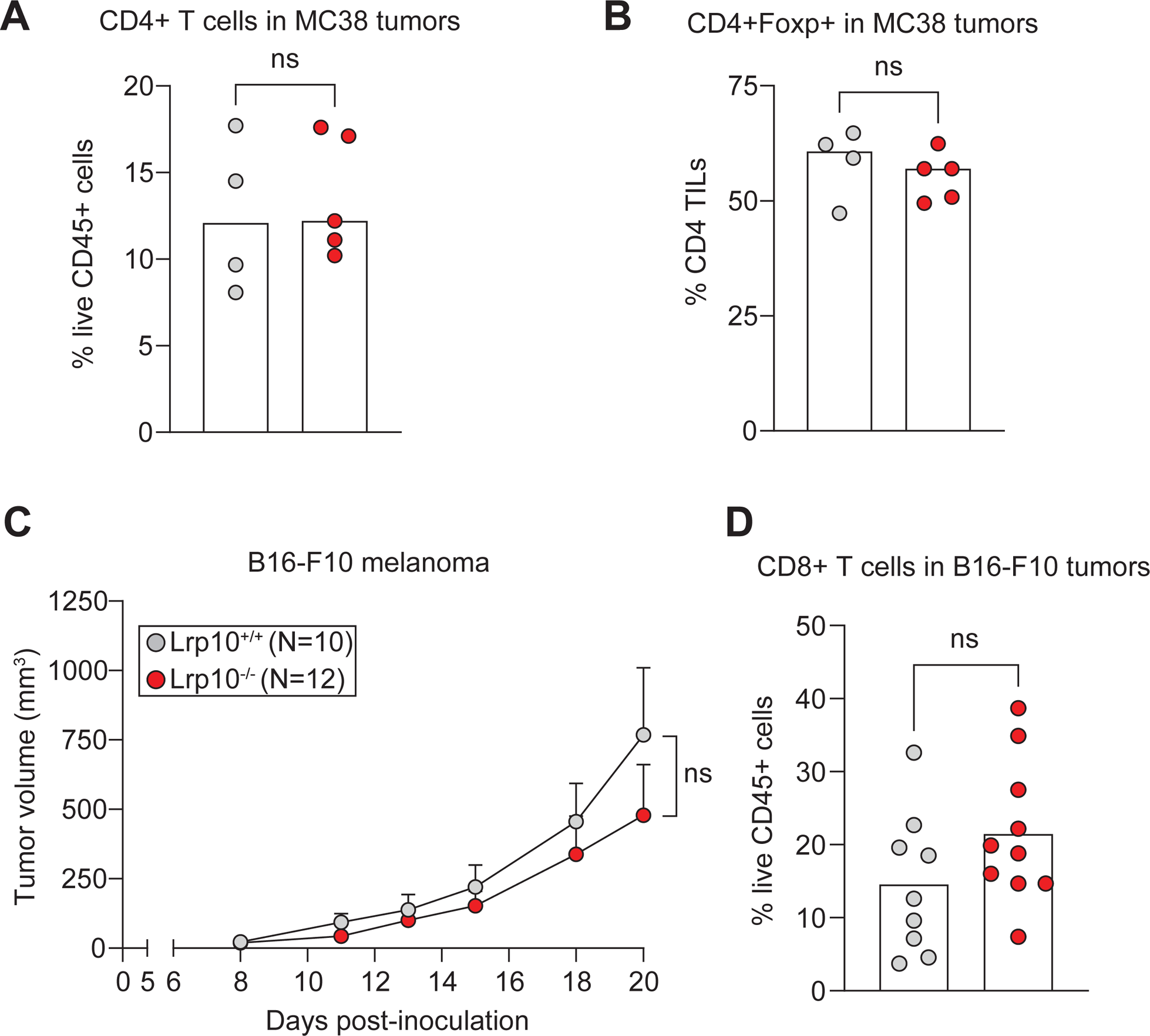
**A, B:** Frequency of CD4 T cells (A) and CD4+Foxp3+ Tregs (B) infiltrating D18 MC38 tumors on *Lrp10^+/+^* and *Lrp10^−/−^* mice. **C:** Mean tumor volumes of B16F10 melanoma cell injected subcutaneously into *Lrp10^+/+^* and *Lrp10^−/−^* mice. Data is presented as mean +/− SEM and representative of three independent experiments. **D:** Frequency of CD8 T cells infiltrating B16F10 tumors on D20. Data is combined from two independent experiments.

**FIGURE S5:**
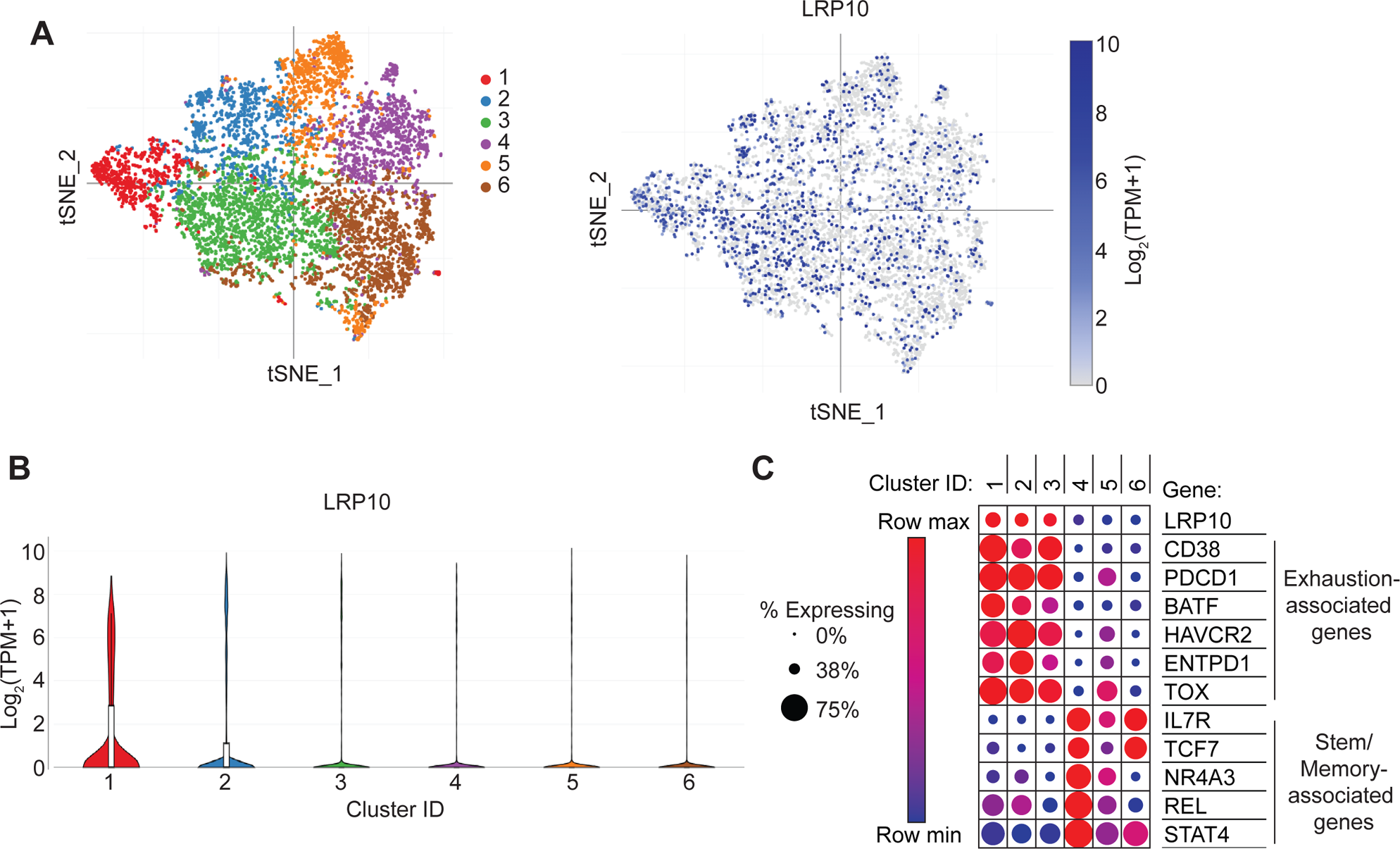
**A:** T-distributed stochastic neighbor embedding (t-SNE) plot of CD8 TILs from melanoma patients from Sade-Feldman, et al. (Sade-Feldman *et al*., 2018) and the distribution of Lrp10 expression. Data was visualized through the Broad Institute Single Cell Portal. Clusters 1-3 were enriched for exhaustion genes and associated with resistance to checkpoint inhibitors while clusters 4-6 were enriched for stem-memory genes and associated with checkpoint inhibitor responsiveness. **B:** Violin plots showing the distribution of Lrp10 in different clusters from (A). **C:** Dot plot showing overlap of Lrp10 expression with known exhaustion markers and negative correlation with stem/memory markers.

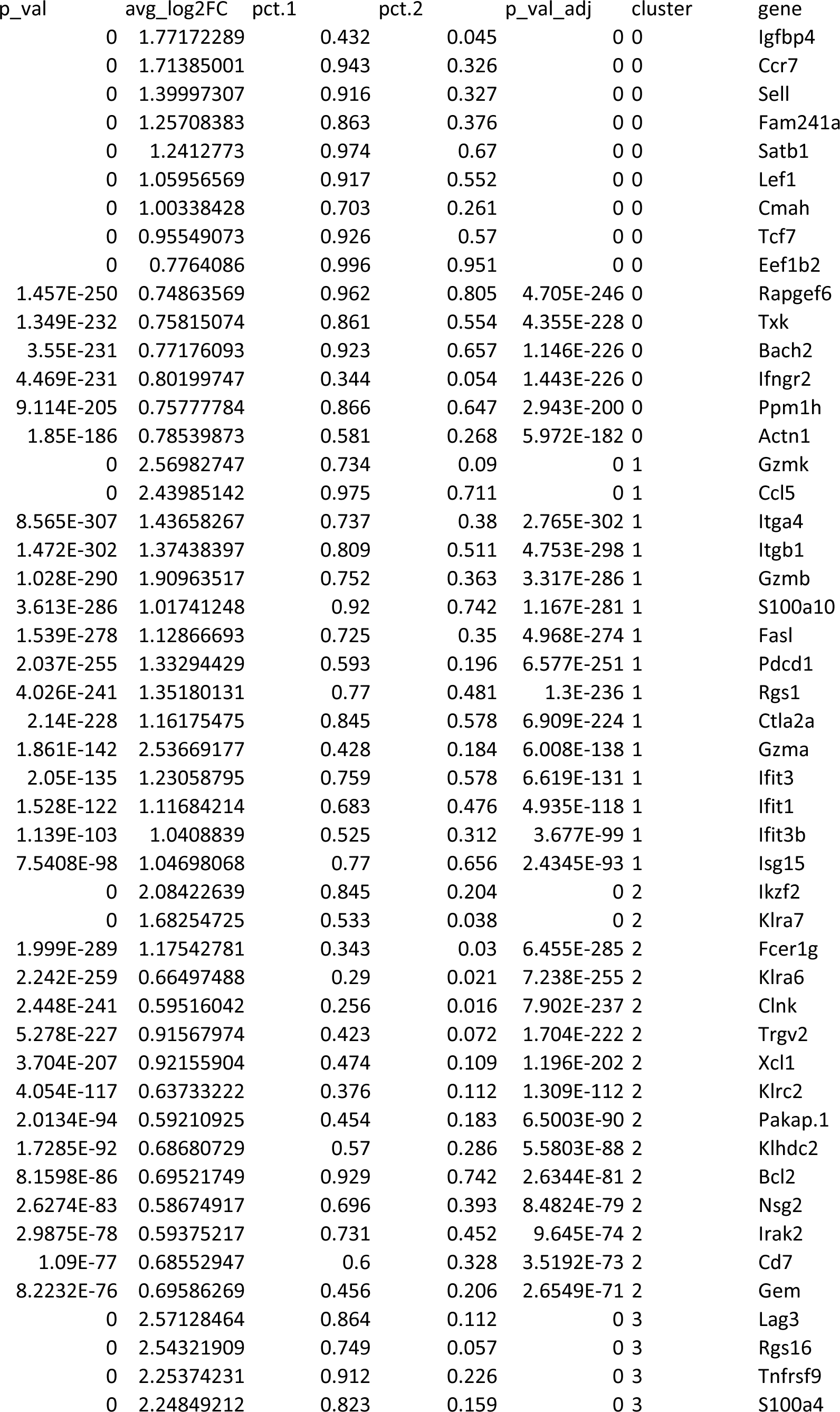
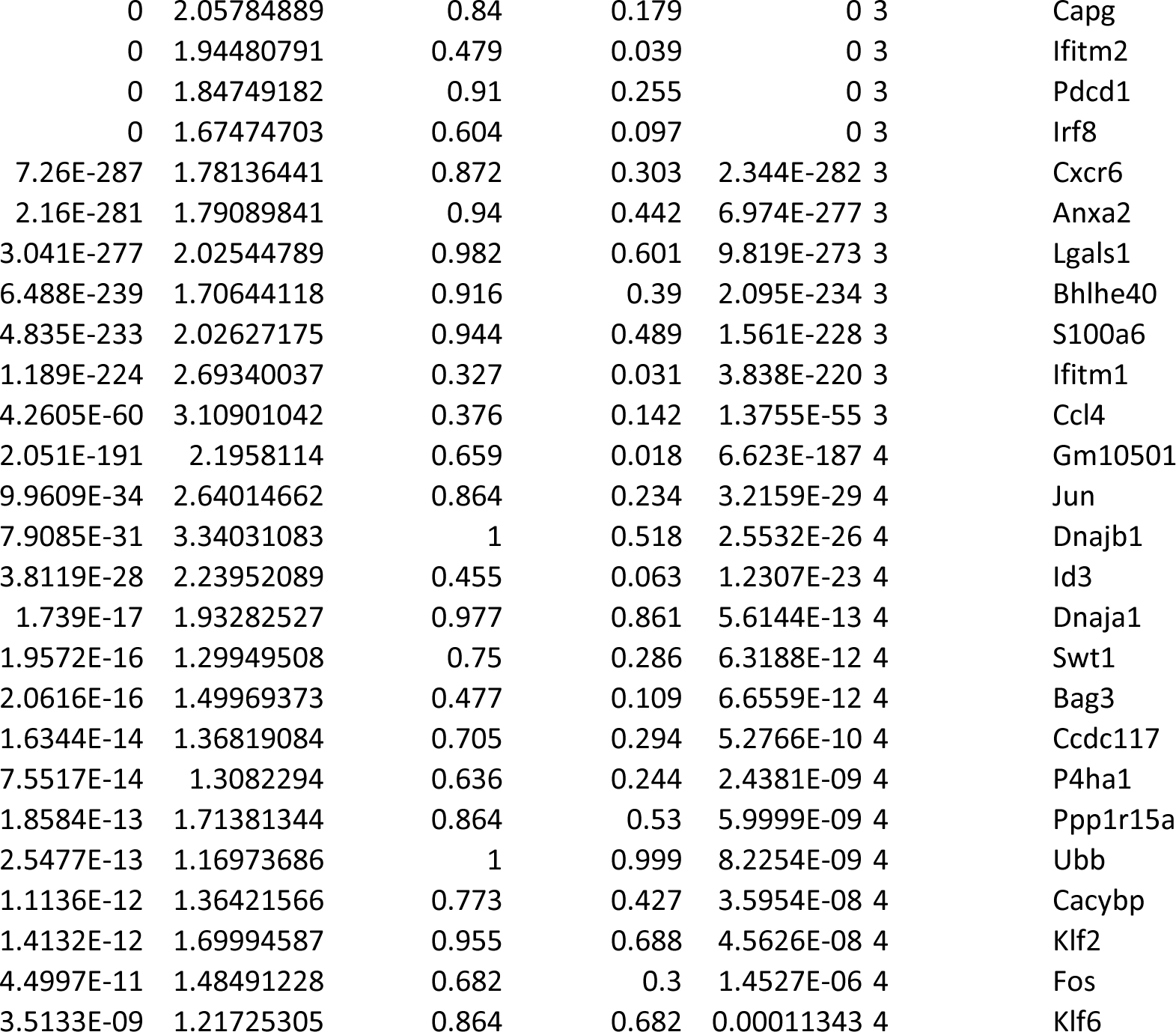

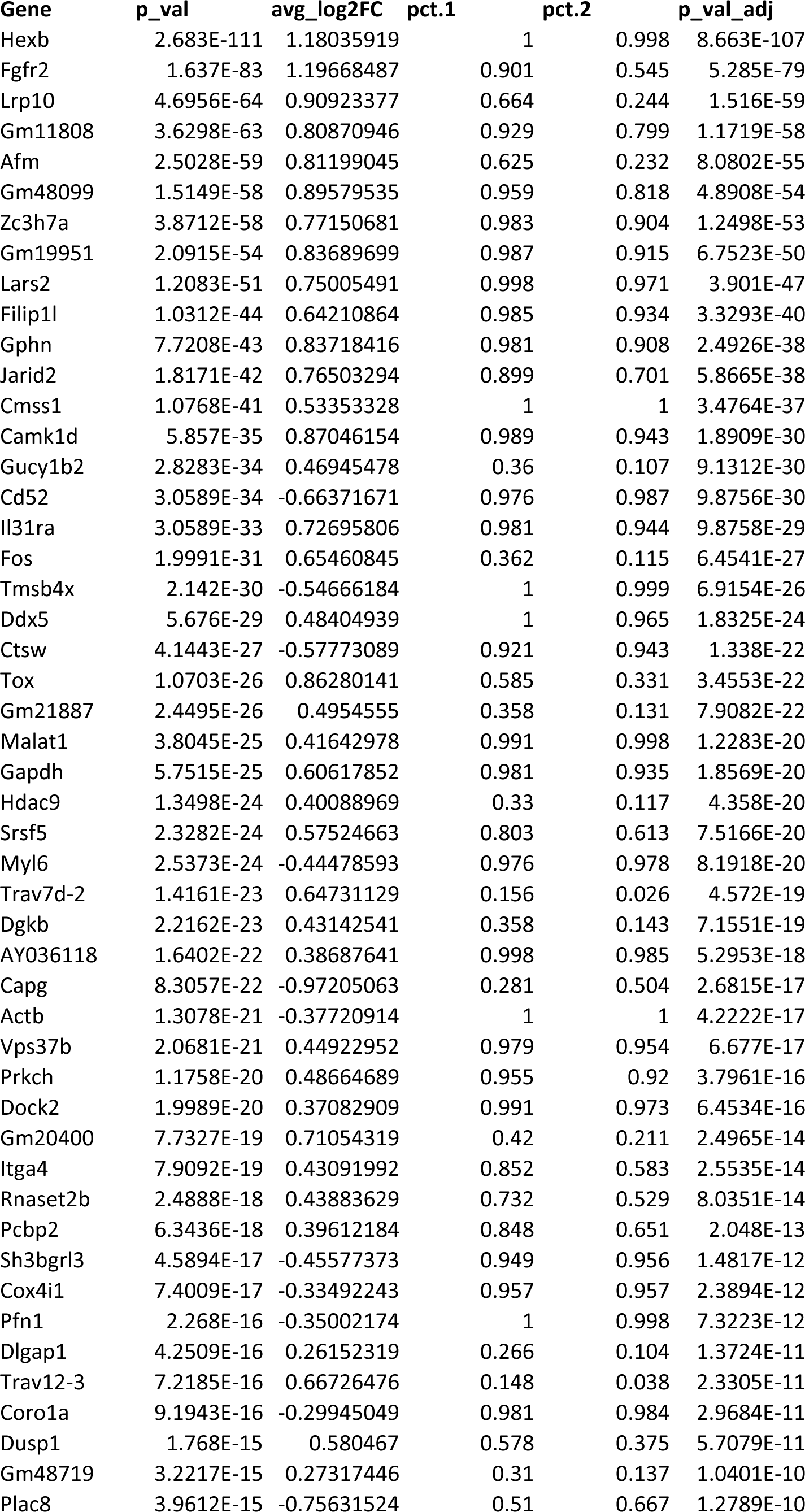
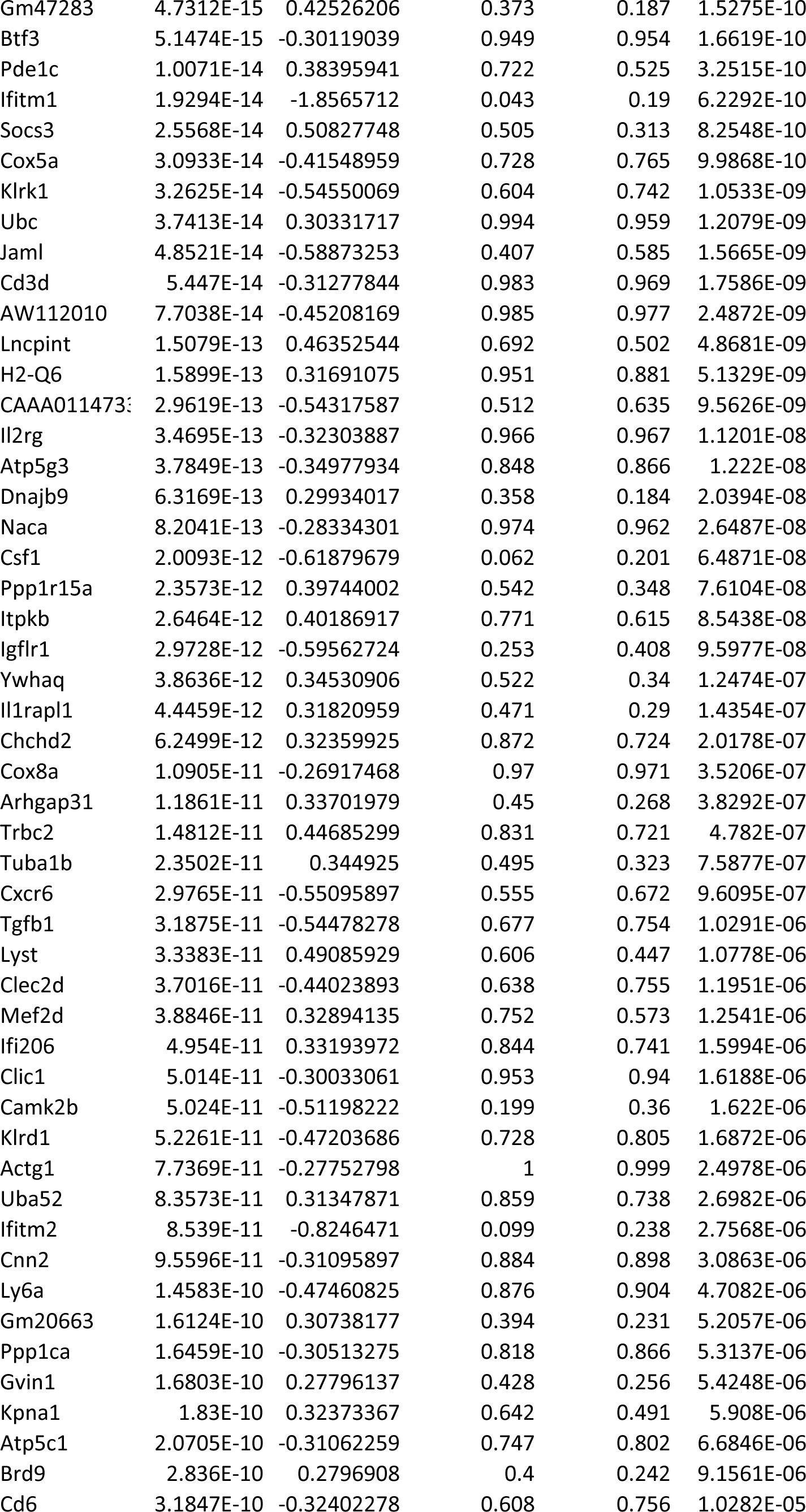
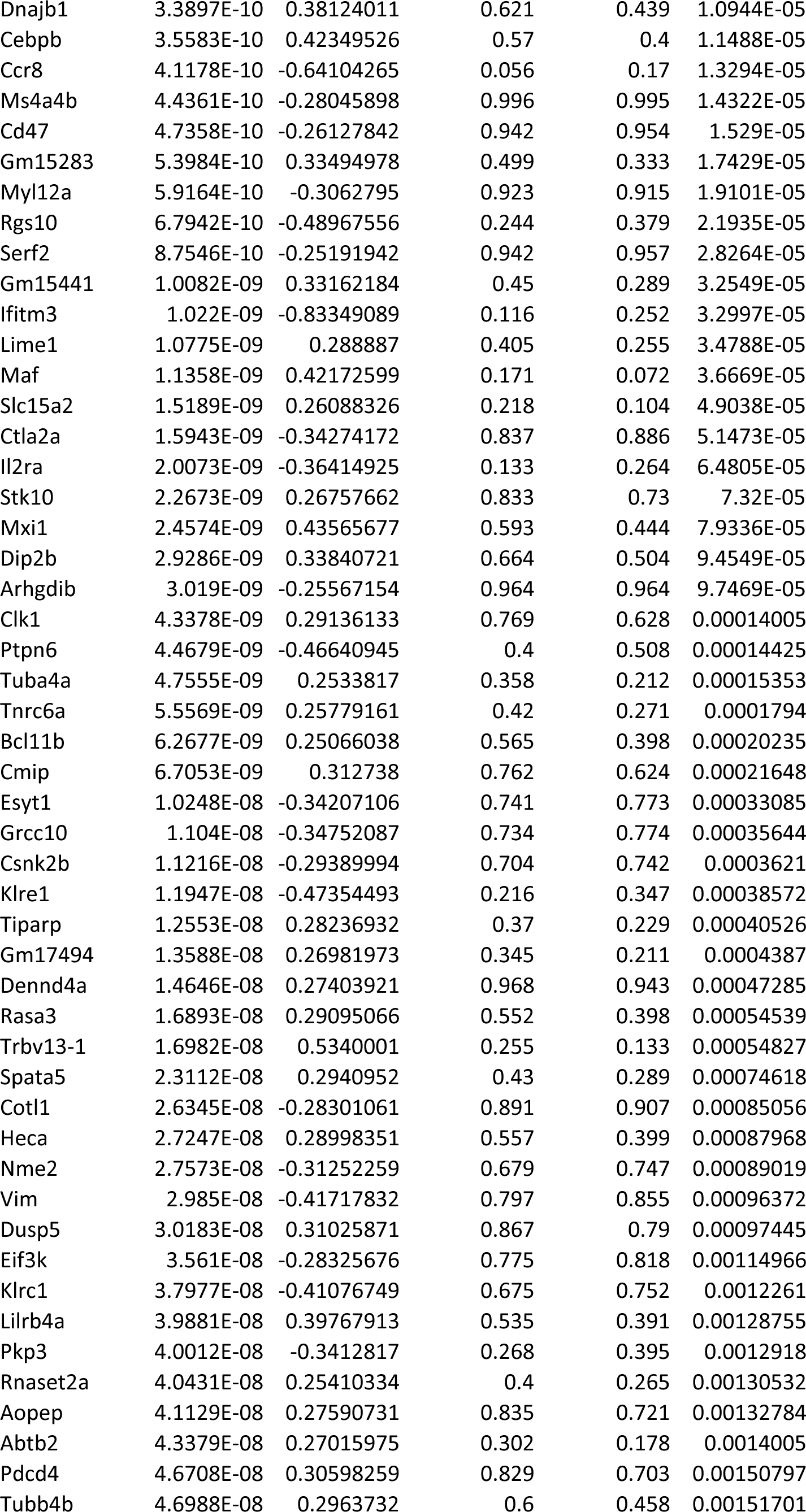
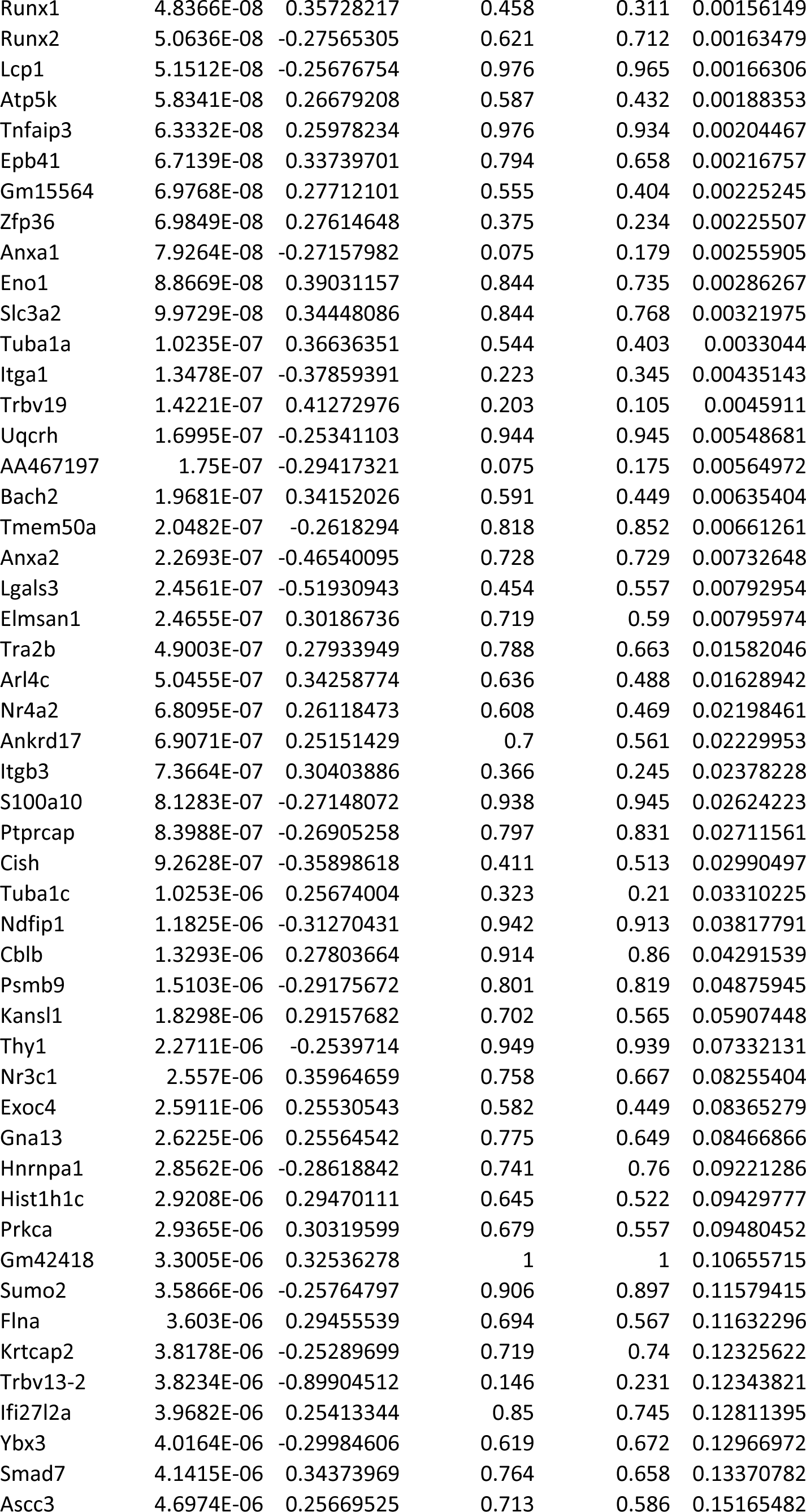
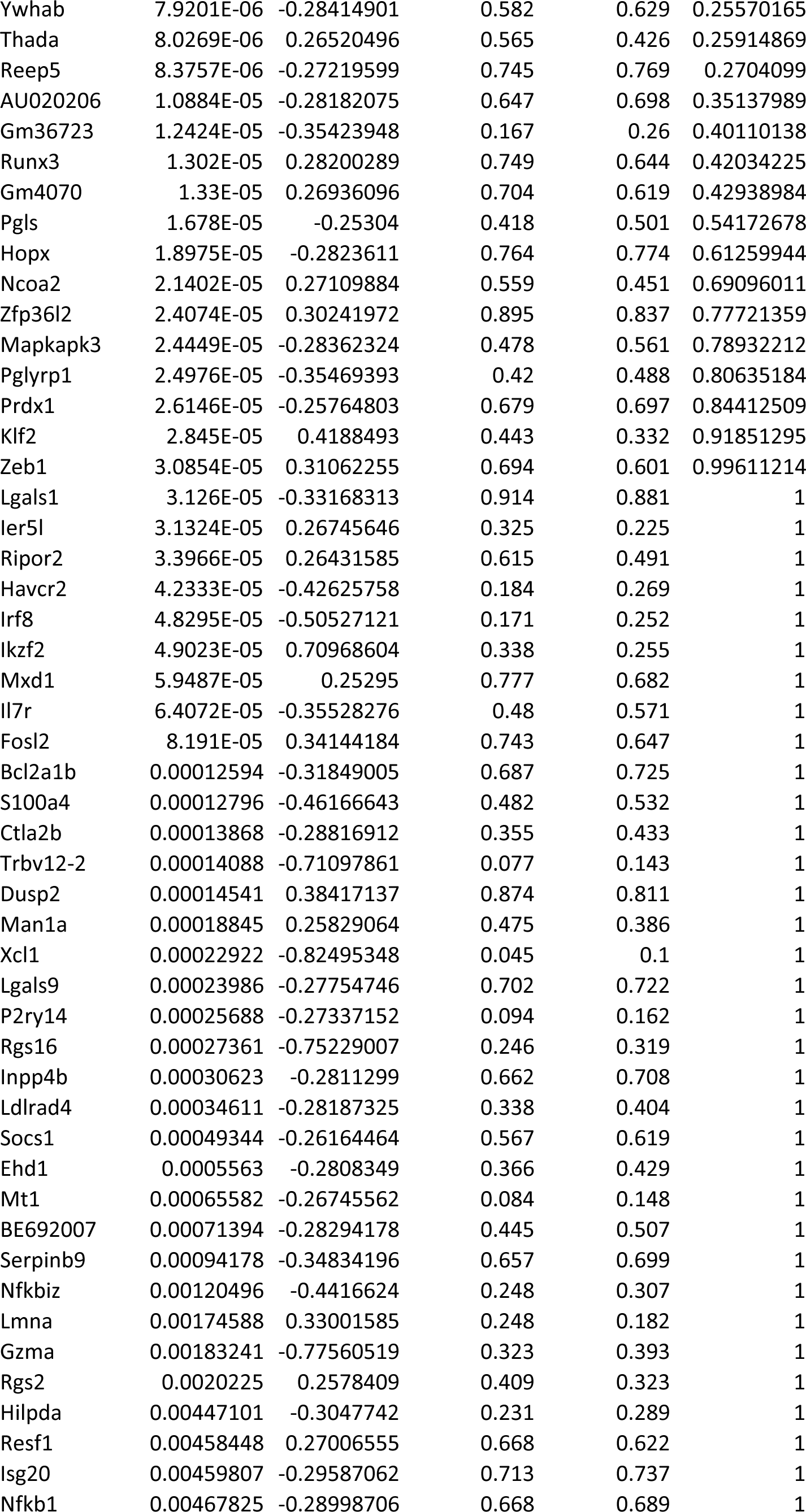
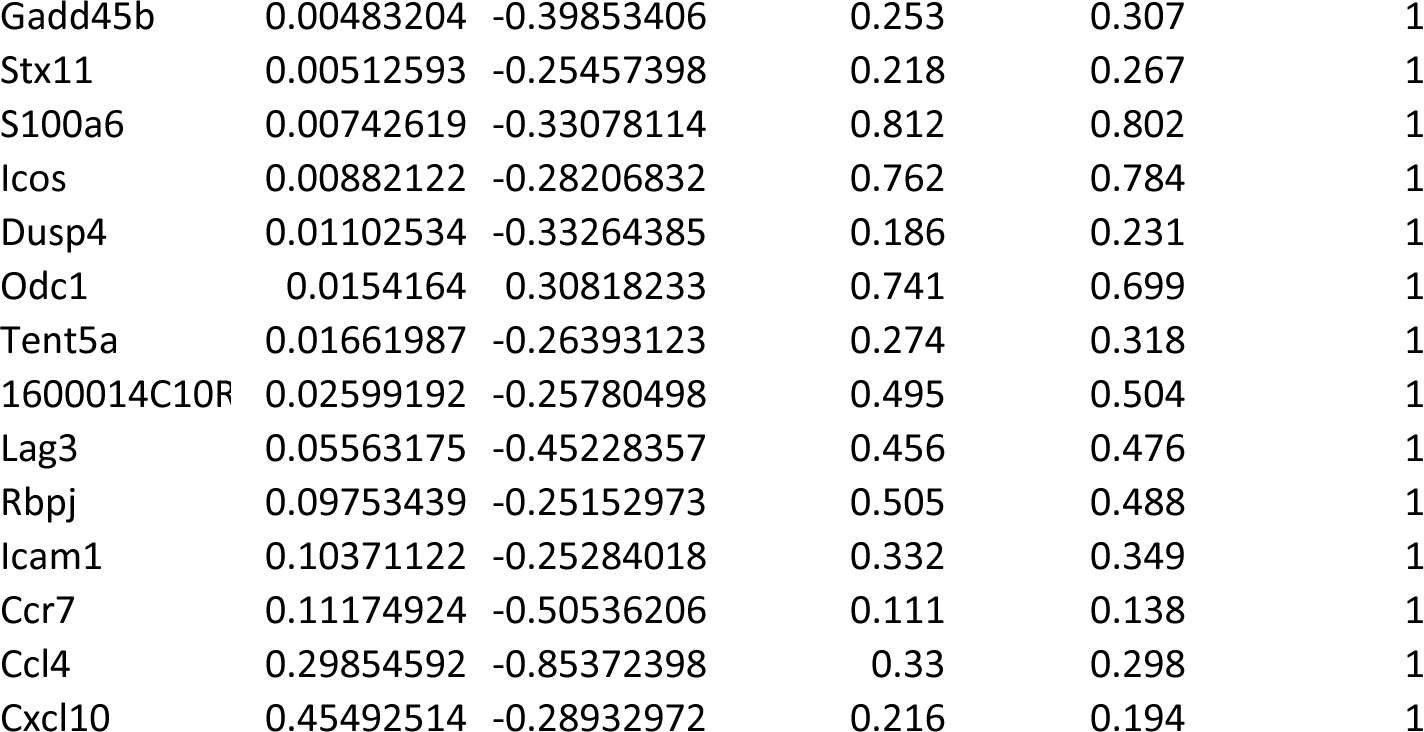

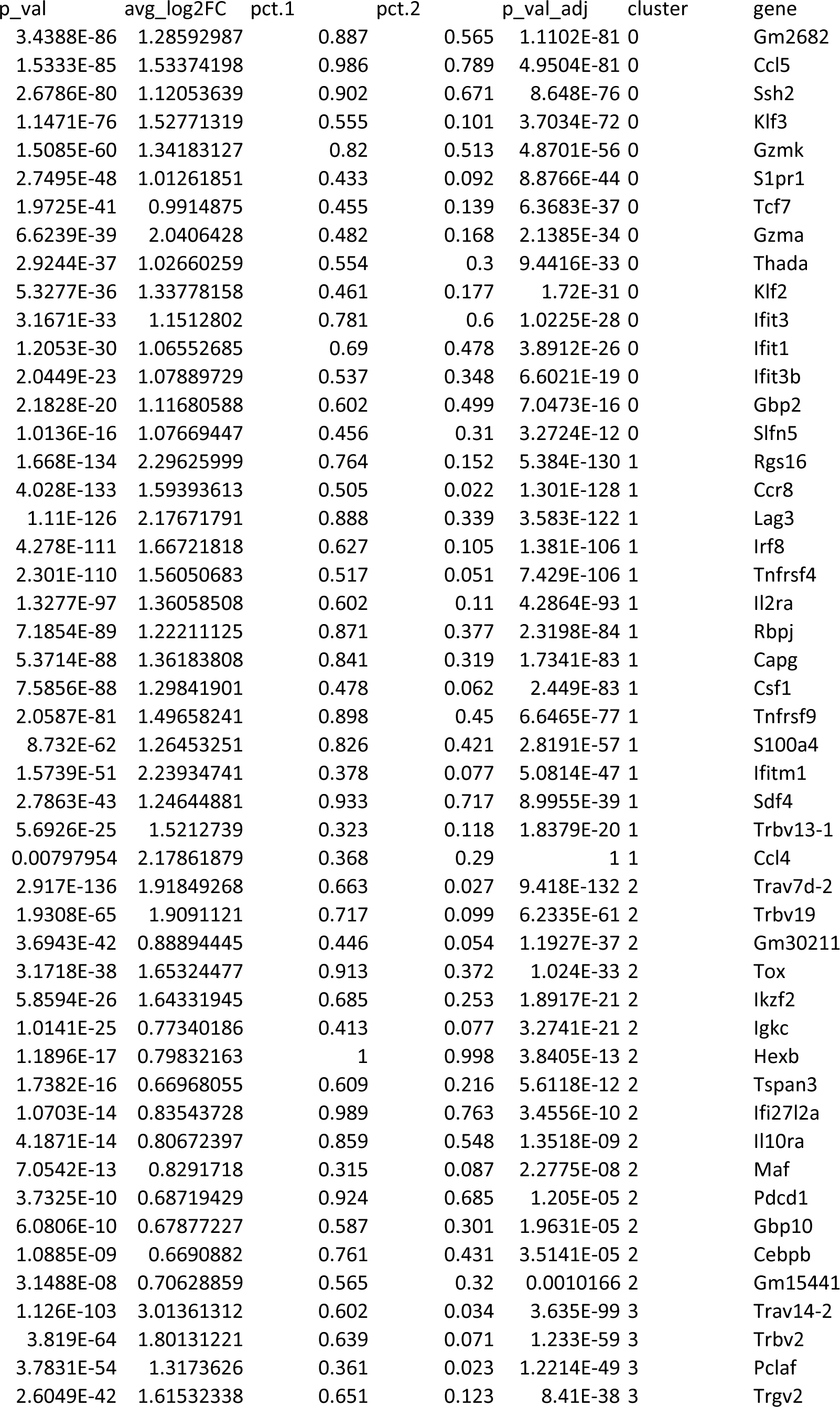
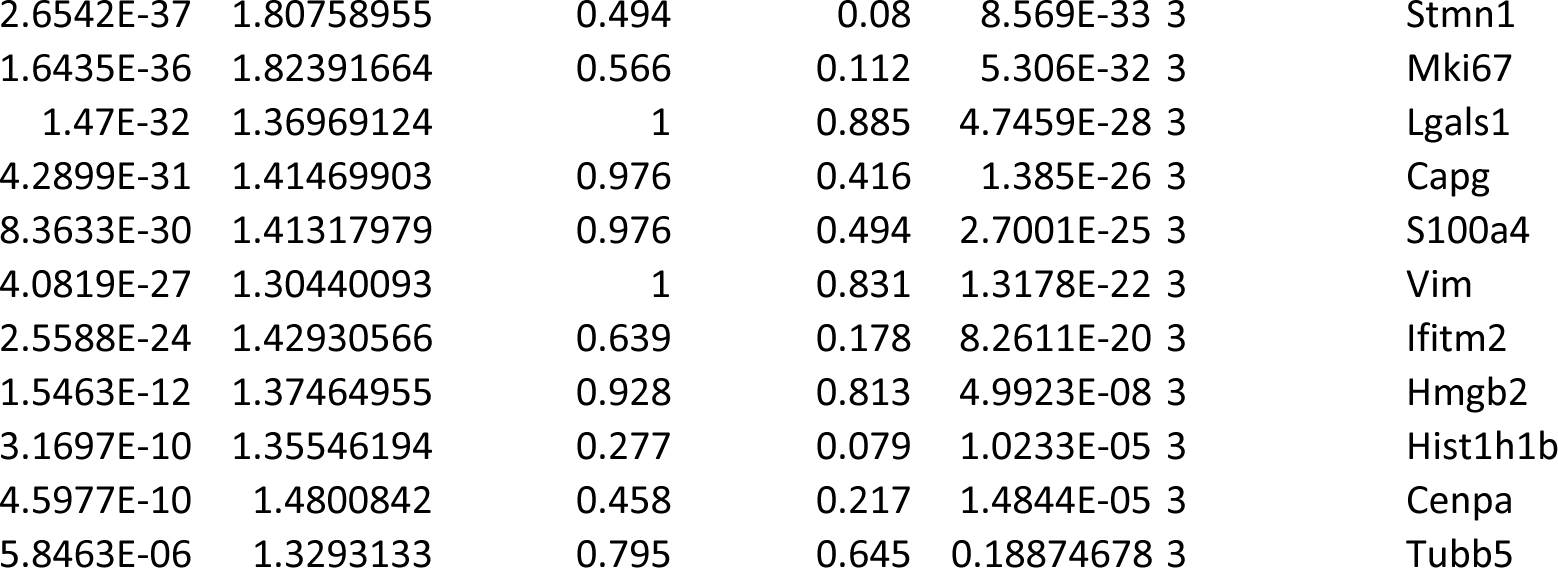

